# Neuronal APOE4 alone is sufficient to drive tau pathology, neurodegeneration, and neuroinflammation in an Alzheimer’s disease mouse model

**DOI:** 10.1101/2025.11.25.690488

**Authors:** Jessica Blumenfeld, Yaqiao Li, Min Joo Kim, Oscar Yip, Luokang Yao, Samuel De Leon, Rebecca Fulthorpe, David Shostak, Zoe Platow, Kaylie Suan, Yanxia Hao, Nicole Koutsodendris, Claire Ellis, Jeremy Nguyen, Yadong Huang

**Affiliations:** Gladstone Institute of Neurological Disease, Gladstone Institutes, San Francisco, CA, USA; Neuroscience Graduate Program, University of California, San Francisco, San Francisco, CA, USA; Gladstone Center for Translational Advancement, Gladstone Institutes, San Francisco, CA, USA; Biomedical Sciences Graduate Program, University of California, San Francisco, San Francisco, CA, USA; Developmental and Stem Cell Biology Graduate Program, University of California, San Francisco, San Francisco, CA, USA; Departments of Pathology and Neurology, University of California, San Francisco, San Francisco, CA, USA

## Abstract

Apolipoprotein E4 (APOE4), the strongest genetic risk factor for late-onset Alzheimer’s disease (AD), exacerbates tau tangles, amyloid plaques, neurodegeneration, and neuroinflammation—the pathological hallmarks of AD. While astrocytes are the primary producers of APOE in the CNS, neurons increase APOE expression under stress and aging. Prior work established that neuronal APOE4 is essential for AD pathogenesis, but whether it is sufficient to drive disease remained unknown. We generated a PS19 tauopathy mouse model selectively expressing APOE4 in neurons. Neuronal APOE4 alone proved sufficient to promote pathological tau accumulation and propagation, neurodegeneration, and neuroinflammation to levels comparable to a tauopathy model with human APOE4 knocked-in globally. Single-nucleus RNA sequencing further revealed similar transcriptomic changes in neurons and glia of both models. Together, these findings demonstrate that neuronal APOE4 alone can initiate and propagate AD pathologies, underscoring its pivotal role in disease pathogenesis and its potential as a therapeutic target.

## Introduction

Alzheimer’s Disease (AD), the most prevalent neurodegenerative disorder, is clinically characterized by a progressive decline in memory and cognition. Pathologically, AD is defined by several cellular phenotypes, including the accumulation of hyperphosphorylated tau (p-tau) and amyloid-β (Aβ) plaques, loss of hippocampal and neuronal mass, and robust neuroinflammation^1–5^.

The greatest genetic risk factor for late-onset AD is apolipoprotein E4 (APOE4)^6,7^. APOE4 is one of three major isoforms of human APOE, alongside APOE2 and APOE3. The most prevalent isoform, APOE3, is present in approximately 75% of the general population and in ∼30% of AD cases^8,9^. In contrast, APOE4, though carried by only ∼24% of the general population^10^, is found in approximately 65% of individuals with AD^5,11^, and is thus disproportionally overrepresented in the AD population.

At the pathological level, APOE4 exacerbates various AD phenotypes. APOE4 is associated with increased p-tau accumulation in both AD brains and model systems^1,12,13^, as well as greater hippocampal atrophy and neuron loss in patient brains and mouse models^12,14–17^. Elevated gliosis and inflammatory cytokines have also been observed with APOE4^12,14,18–20^. Given the involvement of multiple cell types in the central nervous system (CNS) in AD, a key question remains: how does the cellular source of APOE influence disease pathogenesis?

While astrocytes produce the majority of APOE in the CNS, neurons display a striking upregulation of APOE expression under conditions of stress and disease^21–27^. In both APOE knock-in (APOE-KI) mice and human AD brains, neuronal APOE expression closely tracks disease progression, with proportions of neurons that highly express APOE peaking at the start of pathologies during the mild cognitive impairment (MCI) stage^16^, suggesting a potential role of neuronal APOE in initiating pathology. Consistent with this, selective neuronal deletion of APOE4 mitigates key disease features, including hippocampal atrophy and neuronal loss^14,16^, p-tau accumulation, and gliosis^14^, indicating that neuronal APOE4 expression is necessary for AD pathogenesis.

The evidence demonstrating neuronal APOE4 is necessary to drive AD raises an important next question: is neuronal APOE4 alone sufficient to induce the full spectrum of AD pathologies? To address this question, we have generated a novel tauopathy mouse model in which APOE4 expression is restricted only to neurons. Comparison with a global APOE4 knock-in (APOE4-KI) tauopathy model revealed that neuronal APOE4 expression alone was sufficient to recapitulate the key AD hallmarks, including p-tau pathology, hippocampal volume and neuron loss, and neuroinflammation. Together, these findings robustly establish neuronal APOE4 as a central driver of AD pathogenesis.

## Results

### Neuron-specific expression of APOE4 in a tauopathy mouse model

To assess the role of neuronal APOE4 in AD pathogenesis, we utilized a mouse model in which an *APOE4* minigene is inserted under the control of the neuron-specific enolase (NSE) promoter (NSE-E4)^28–33^ (Extended Data Fig. 1a), which has previously been shown to drive robust, pan-neuronal expression of various fusion genes^34,35^. The *APOE4* minigene construct contains the full coding sequence of APOE4, as previously reported (Extended Data Fig. 1a)^28,29^. NSE-E4 mice were crossbred onto a mouse ApoE knockout background to eliminate other cellular sources of APOE. To examine the effects of neuronal APOE4 in a tauopathy context, we further crossbred the NSE-E4 mice with the PS19 tauopathy mouse model, which expresses the human P301S tau mutation^36^. The resulting mice are referred to as PS19/NSE-E4. PS19 mice harboring human *APOE4* or *APOE3* knock-in alleles (referred to as PS19/E4 and PS19/E3) served as diseased and healthy controls, respectively. Mice were analyzed at 10 months of age to capture advanced stages of PS19 pathology.

To confirm neuron-specific expression of APOE in the PS19/NSE-E4 model, we performed immunostaining for APOE in the hippocampus alongside markers for the four major CNS cell types: neurons (NeuN), astrocytes (GFAP), microglia (IBA1), and oligodendrocytes (Olig2). In PS19/NSE-E4 mice, APOE colocalized with neurons, but not with astrocytes, microglia, or oligodendrocytes (Extended Data Fig. 1b,c). By contrast, PS19/E4 mice show prominent APOE colocalization within astrocytes, with additional signal observed within microglia and neurons as well (Extended Data Fig. 1b,c). Altogether, these findings recapitulate previous reports of the original NSE-E4 line^28–33^ and confirm that the PS19/NSE-E4 model achieves neuron-specific APOE expression.

### Neuronal APOE4 alone is sufficient to promote tau pathology

Given the tauopathy background of this line, we first investigated the impact of neuron-specific APOE4 expression on the accumulation of hyperphosphorylated tau (p-tau). Immunostaining using a p-tau-specific antibody, AT8, we assessed whether expression of neuronal APOE4 alone would induce p-tau accumulation. Strikingly, PS19/NSE-E4 mice exhibited markedly greater hippocampal p-tau burden than PS19/E3 mice, showing an approximately 77% increase in p-tau coverage area (Fig. 1a,b). Notably, PS19/NSE-E4 mice displayed p-tau burden comparable to that of PS19/E4 mice (∼13% increase) (Fig. 1a,b).

**Fig. 1.**
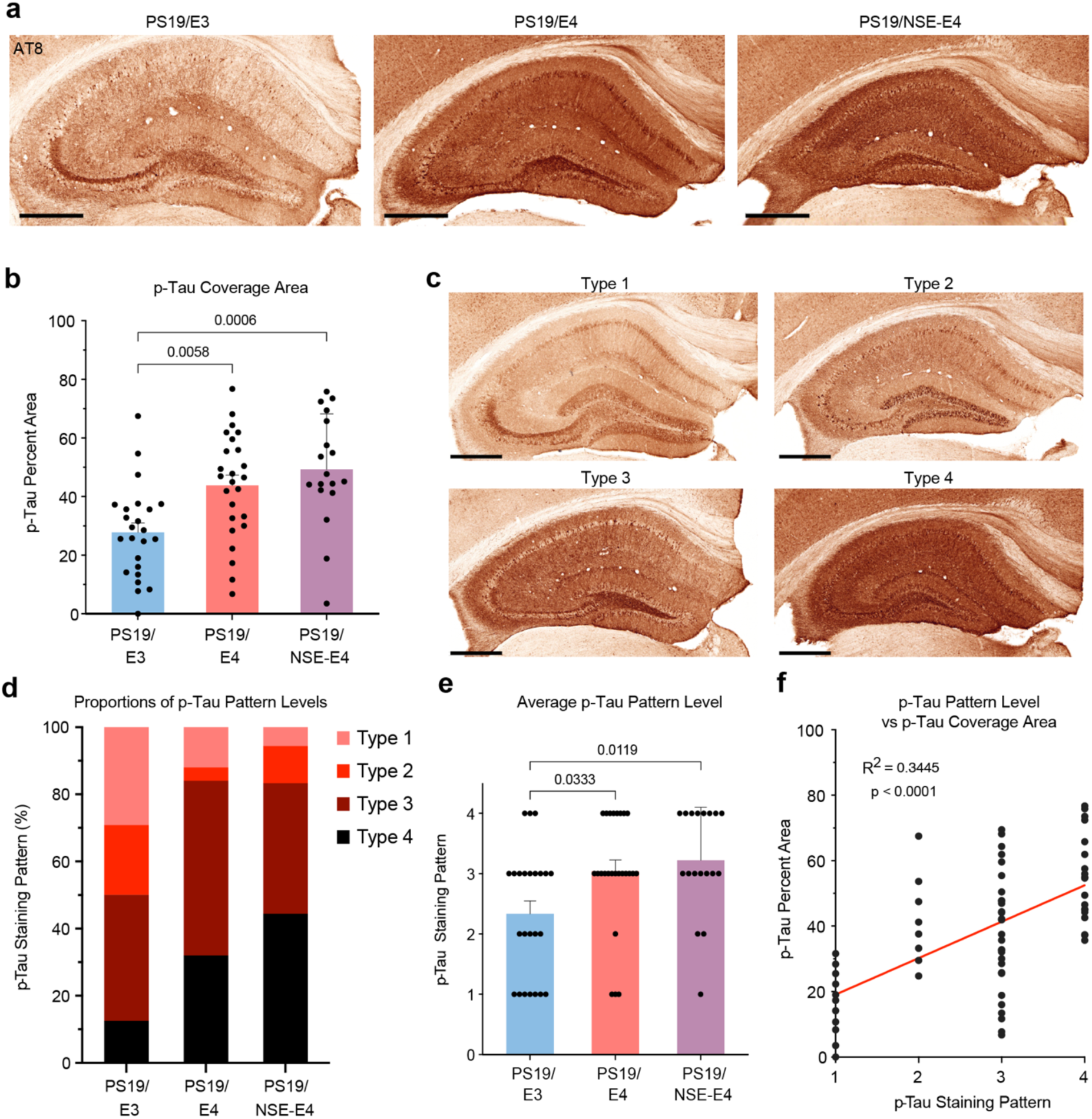
Neuronal APOE4 alone exacerbates Tau pathology in tauopathy mice. **a**, Representative images of the hippocampus from 10-month-old PS19/E3, PS19/E4, and PS19/NSE-E4 mice, stained with p-Tau-specific monoclonal AT8 antibody (scale bar, 500 µm). **b**, Quantifications of percent p-Tau coverage of the hippocampus of PS19/E3, PS19/E4, and PS19/NSE-E4 mice. **c**, Representative images of the four levels of AT8 hippocampal staining patterns (scale bar, 500 µm). **d**, Stacked bar graph of the distribution of p-Tau staining patterns across PS19/E3, PS19/E4, and PS19/NSE-E4 mice. **e**, Quantification of p-Tau staining pattern severity across PS19/E3, PS19/E4, and PS19/NSE-E4 mice. **f**, Correlation of hippocampal p-Tau staining pattern and percent p-Tau coverage area. For quantifications in **b**,**d**,**e**,**f**, PS19/E3 *n* = 24; PS19/E4 *n* = 25; and PS19/NSE-E4 *n* = 18. In **b**,**e**,**f**, data is expressed as mean ± s.e.m and was assessed via one-way analysis of variance (ANOVA) with Tukey’s post hoc multiple comparisons test.

Previous studies have characterized four major p-tau staining patterns in PS19/APOE-KI models, corresponding to progressively diseased states (Fig. 1c)^12,15,37^. In type 1, p-tau is primarily localized to the mossy fibers and diffusely spread throughout the dentate gyrus (DG) granule layer and CA1 pyramidal layer. Type 2 features dense tau tangles within the cell bodies of DG granule cells and CA3 pyramidal neurons. In type 3, p-tau extends into the dendrites of pyramidal neurons, producing prominent staining in the stratum radiatum. Type 4 represents the most advanced stage, with diffuse, grainy staining appearing throughout the entirety of the hippocampus. To further evaluate the effect of neuronal APOE4 on tau pathology, we classified hippocampal p-tau patterns for each mouse (Fig. 1c).

Approximately 50% of PS19/E3 mice exhibited the less severe type 1 or type 2 p-tau patterns (Fig. 1d). In contrast, only ∼16% of PS19/E4 and PS19/NSE-E4 mice fell into these categories, with both groups being largely comprised of type 3 and type 4 p-tau staining (Fig. 1d). While both PS19/E4 and PS19/NSE-E4 mice exhibited a more advanced average p-tau pattern type than PS19/E3 mice (Fig. 1e), PS19/NSE-E4 mice displayed a higher proportion in type 4 than PS19/E4 mice. Whereas 32% of PS19/E4 mice displayed type 4 p-tau pathology, about 44% were type 4 in the PS19/NSE-E4 group (Fig. 1d), suggesting that increased neuronal APOE4 expression can further drive tauopathy progression. Notably, tau staining type and percent p-tau coverage area correlated, indicating that hippocampal coverage reflects p-tau progression (Fig. 1f). Together, these findings demonstrate that neuron-specific expression of APOE4 alone is sufficient to induce and exacerbate tau pathology.

### Neuronal APOE alone is sufficient to drive neurodegeneration

We next investigated whether selective neuronal expression of APOE4 could promote neurodegeneration. Previous studies have shown that both global and neuron-specific APOE deletion mitigates hippocampal atrophy^12,14^. To determine whether neuronal APOE4 expression alone could drive degeneration, we measured hippocampal volume at 10 months of age. PS19/NSE-E4 mice exhibited pronounced atrophy, with hippocampal volumes comparable to those observed in PS19/E4 mice and nearly 40% smaller than the hippocampi of PS19/E3 controls (Fig. 2a,b).

**Fig. 2.**
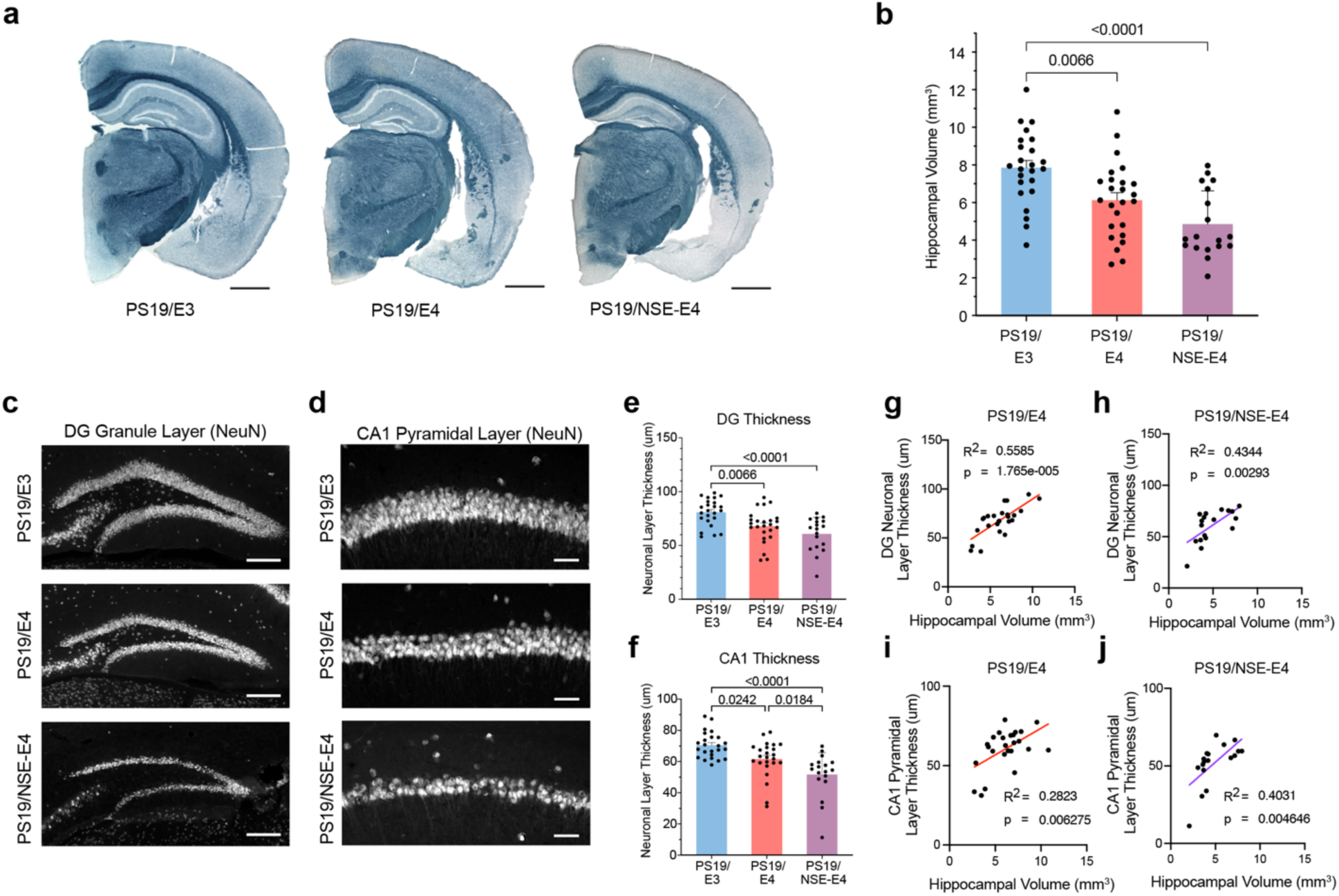
Neuronal APOE4 alone potentiates neurodegeneration and hippocampal atrophy in tauopathy mice. **a**, Representative images of cross-sectional hippocampal brain tissue from 10-month-old PS19/E3, PS19/E4, and PS19/NSE-E4 mice, incubated in Sudan Black to enhance hippocampal visibility (scale bar, 1000 µm). **b**, Quantifications of hippocampal volume of PS19/E3, PS19/E4, and PS19/NSE-E4 mice. **c**,**d**, Representative images of the DG granule layer (scale bar, 200 µm) (**c**) and the CA1 pyramidal layer subfield (scale bar, 50 µm) (**d**) of PS19/E3, PS19/E4, and PS19/NSE-E4 mice, labeled with NeuN to capture neurons. **e**,**f**, Quantification of DG granule layer thickness (**e**) and CA1 pyramidal layer (**f**) of PS19/E3, PS19/E4, and PS19/NSE-E4 mice. **g**,**h**, Correlation of hippocampal volume and DG layer thickness in PS19/E4 mice (**g**) and PS19/NSE-E4 mice (**h**). **i**,**j**, Correlation of hippocampal volume and CA1 layer thickness in PS19/E4 mice (**i**) and PS19/NSE-E4 mice (**j**). For quantifications throughout, PS19/E3 *n* = 24; PS19/E4 *n* = 25; and PS19/NSE-E4 *n* = 18. In **b**,**e**,**f**, data is expressed as mean ± s.e.m and was assessed via one-way analysis of variance (ANOVA) with Tukey’s post hoc multiple comparisons test. Pearson correlation analysis was used to determine significance in **g**,**h**,**i**,**j**.

To assess degeneration at the cellular level, we also quantified neuronal layer thickness following neuronal (NeuN) immunolabeling. Both PS19/E4 and PS19/NSE-E4 mice showed significant neuron loss relative to PS19/E3 controls in the DG granule layer and CA1 pyramidal layer (Fig. 2c-f). Strikingly, in the CA1 region, PS19/NSE-E4 mice exhibited even greater neuronal layer thinning than PS19/E4 mice (Fig. 2d,f). Given that hippocampal p-tau accumulation and subsequent atrophy have been found to begin in the CA1 region in AD patients^38,39^, this region-specific vulnerability is consistent with disease progression in humans.

Notably, degeneration of the DG granule and CA1 pyramidal layers strongly correlated with hippocampal atrophy in both PS19/E4 and PS19/NSE-E4 mice (Fig. 2g-j), suggesting that neuronal loss in these regions accounts for much of the observed hippocampal volume reduction. Altogether, these data demonstrate that neuronal APOE4 alone is sufficient to induce neurodegeneration at both cellular and hippocampal levels.

### Neuronal APOE4 alone is sufficient to promote gliosis

Given that neuroinflammation is a well-characterized feature of PS19/E4 mice^12,14,15,37^, we next examined how selective neuronal expression of APOE4 influences glial activation. We began with quantifying microgliosis in the hippocampus of 10-month-old mice. Both PS19/NSE-E4 and PS19/E4 mice exhibited markedly elevated levels of IBA1-positive microglia coverage compared to PS19/E3 controls (Fig. 3a,b). Additionally, CD68-positive immunolabeling, used to identify activated and phagocytic microglia, was also significantly increased in PS19/NSE-E4 and PS19/E4 mice relative to PS19/E3 controls (Fig. 3c,d). Intriguingly, despite only having neuronal APOE4 (and therefore lacking microglial APOE4), the PS19/NSE-E4 mice displayed gliosis levels comparable to PS19/E4 mice, suggesting that neuronal APOE4 alone is sufficient to elicit inflammatory microglial responses. While APOE upregulation is a known feature of disease-associated microglia (DAM)^40^, this may indicate that alternative microglial inflammatory pathways are activated downstream of neuronal APOE4 signaling.

**Fig. 3.**
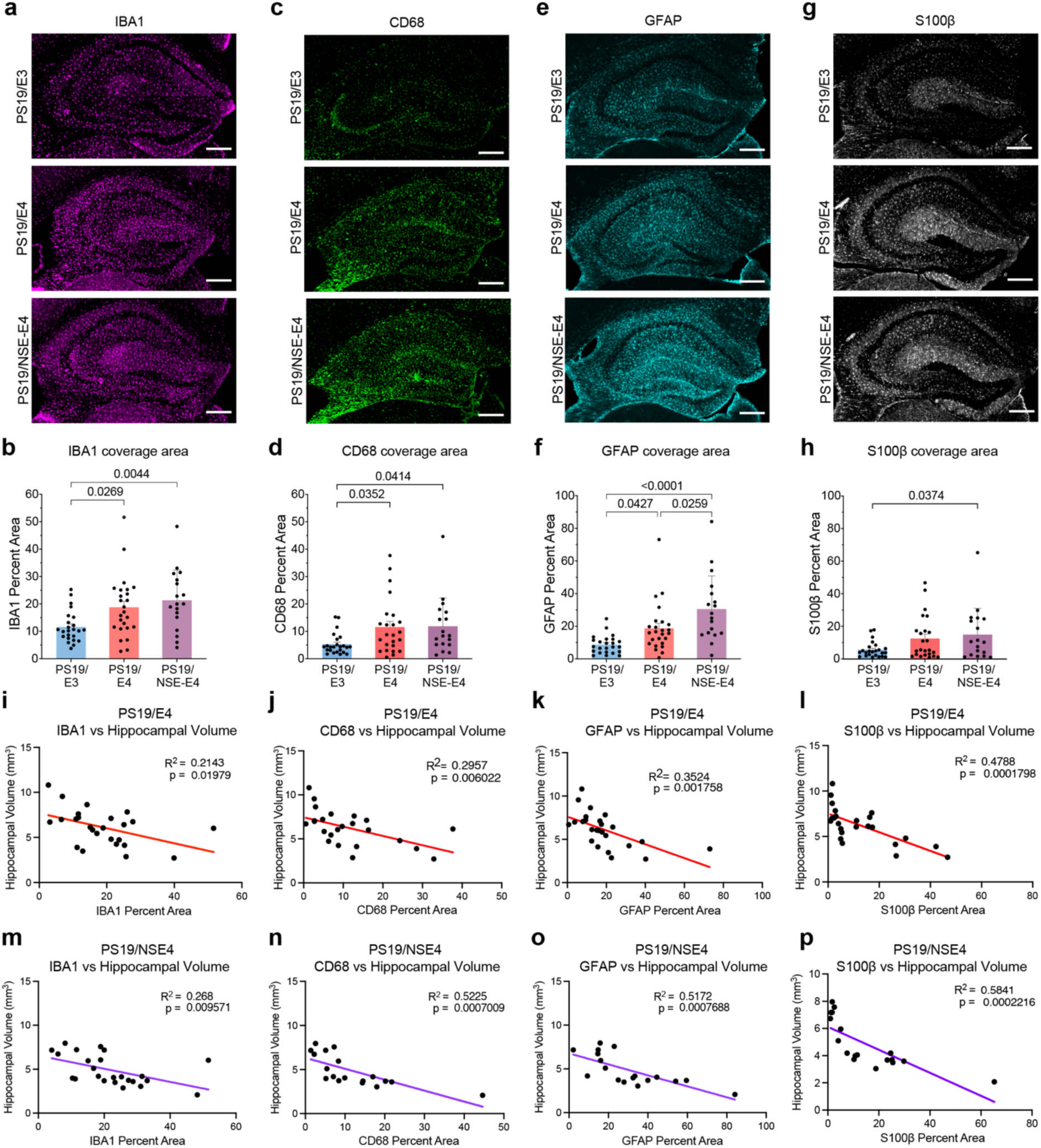
Neuronal APOE4 alone drives gliosis in tauopathy mice. **a**, Representative images of hippocampal brain tissue from 10-month-old PS19/E3, PS19/E4, and PS19/NSE-E4 mice, immunolabeled with microglial marker IBA1. **b**, Quantifications of percent coverage area of IBA1. **c**, Representative images of hippocampal brain tissue from PS19/E3, PS19/E4, and PS19/NSE-E4 mice, immunolabeled with activated microglial marker CD68. **d**, Quantifications of percent coverage area of CD68. **e**, Representative images of hippocampal brain tissue from PS19/E3, PS19/E4, and PS19/NSE-E4 mice, immunolabeled with astrocytic marker GFAP. **f**, Quantifications of percent coverage area of GFAP. **g**, Representative images of hippocampal brain tissue from PS19/E3, PS19/E4, and PS19/NSE-E4 mice, immunolabeled with activated astrocytic marker S100β. **h**, Quantifications of percent coverage area of S100β. In **a,c,e,g,** Scale bars at 300 µm. **i-l**, Correlation of hippocampal volume and IBA1 (**i**), GFAP (**j**), CD68 (**k**), and S100β (**l**) in PS19/E4 mice (*n* = 25). **m-p**, Correlation of hippocampal volume and IBA1 (**m**), GFAP (**n**), CD68 (**o**), and S100β (**p**) in PS19/NSE-E4 mice (*n* = 18). Throughout **b**,**d**,**f**,**h**, PS19/E3, *n* = 24; PS19/E4, *n* = 25; and PS19/NSE-E4, *n* = 18; data is expressed as mean ± s.e.m and was assessed via one-way analysis of variance (ANOVA) with Tukey’s post hoc multiple comparisons test. Pearson correlation analysis was used to determine significance in **i**-**p**.

We next assessed astrocytic activation using GFAP and S100β immunostaining to label total and reactive astrocytes, respectively. Both PS19/E4 and PS19/NSE-E4 mice showed a significant increase in GFAP hippocampal coverage compared with PS19/E3 controls (Fig. 3e,f). Interestingly, PS19/NSE-E4 mice displayed even greater levels of GFAP coverage than PS19/E4 mice, suggesting that neuronal APOE4 specifically may more sensitively potentiate astrocytic inflammatory responses. Similarly, S100β-positive astrocyte coverage increased in PS19/NSE-E4 mice relative to PS19/E3 controls (Fig. 3g,h). Although S100β levels in PS19/E4 mice did not reach statistical significance, they trended higher than PS19/E3 levels and were comparable on average to those in PS19/NSE-E4 mice (Fig. 3g,h). Western blotting on hippocampal lysates also confirmed elevated microgliosis (IBA1) and astrogliosis (GFAP) in PS19/NSE-E4 compared to PS19/E3 mice (Extended Data Fig. 2a-d), further demonstrating neuronal APOE4-driven gliosis.

Across PS19/E4 and PS19/NSE-E4 mice, IBA1, CD68, GFAP, and S100β gliosis levels each correlated strongly with hippocampal atrophy and neuronal thinning in the DG and CA1 (Fig. 3i-p, Extended Data Fig. 2e-t). Notably, the activated glial markers (CD68 and S100β) exhibited even stronger correlations with neurodegeneration than the general glial markers (IBA1 and GFAP), suggesting that activated glia may be directly contributing to degeneration. Together, these findings show that neuronal APOE4 alone is sufficient to elicit robust neuroinflammatory responses comparable to those observed when APOE4 is expressed across all brain cell types.

### Neuronal APOE4 alone is sufficient to drive intracellular HMGB1 translocation

To further investigate whether neuronal APOE4 alone is sufficient to elicit neuroinflammatory responses, we examined neuronal translocation of High Mobility Group Box 1 (HMGB1) across hippocampal subregions. HMGB1 is a nuclear protein that, upon translocation into the cytosol and subsequent release into the extracellular space, functions as a damage-associated molecular pattern (DAMP) that activates inflammatory responses^41–44^. Aberrant HMGB1 release has been implicated in various neurodegenerative disorders^45–48^, and recent work has shown that APOE4 promotes HMGB1 translocation in the PS19 tauopathy model^18^. Although selective deletion of neuronal APOE4 has been shown to attenuate HMGB1 translocation^18^, whether neuronal APOE4 alone is sufficient to drive this process remained unknown.

In both the CA1 pyramidal and the DG granule layers, PS19/NSE-E4 and PS19/E4 mice exhibited significantly higher levels of extranuclear HMGB1 relative to PS19/E3 controls (Fig. 4a-d). Notably, in the DG, PS19/NSE-E4 mice showed even greater HMGB1 translocation than PS19/E4 mice (Fig. 4c,d), suggesting that neuronal APOE4 may be directly involved in neuronal HMGB1 release. These data indicate that neuronal APOE4 expression alone is sufficient to promote HMGB1 release, likely inducing subsequent inflammatory glial response.

**Fig. 4.**
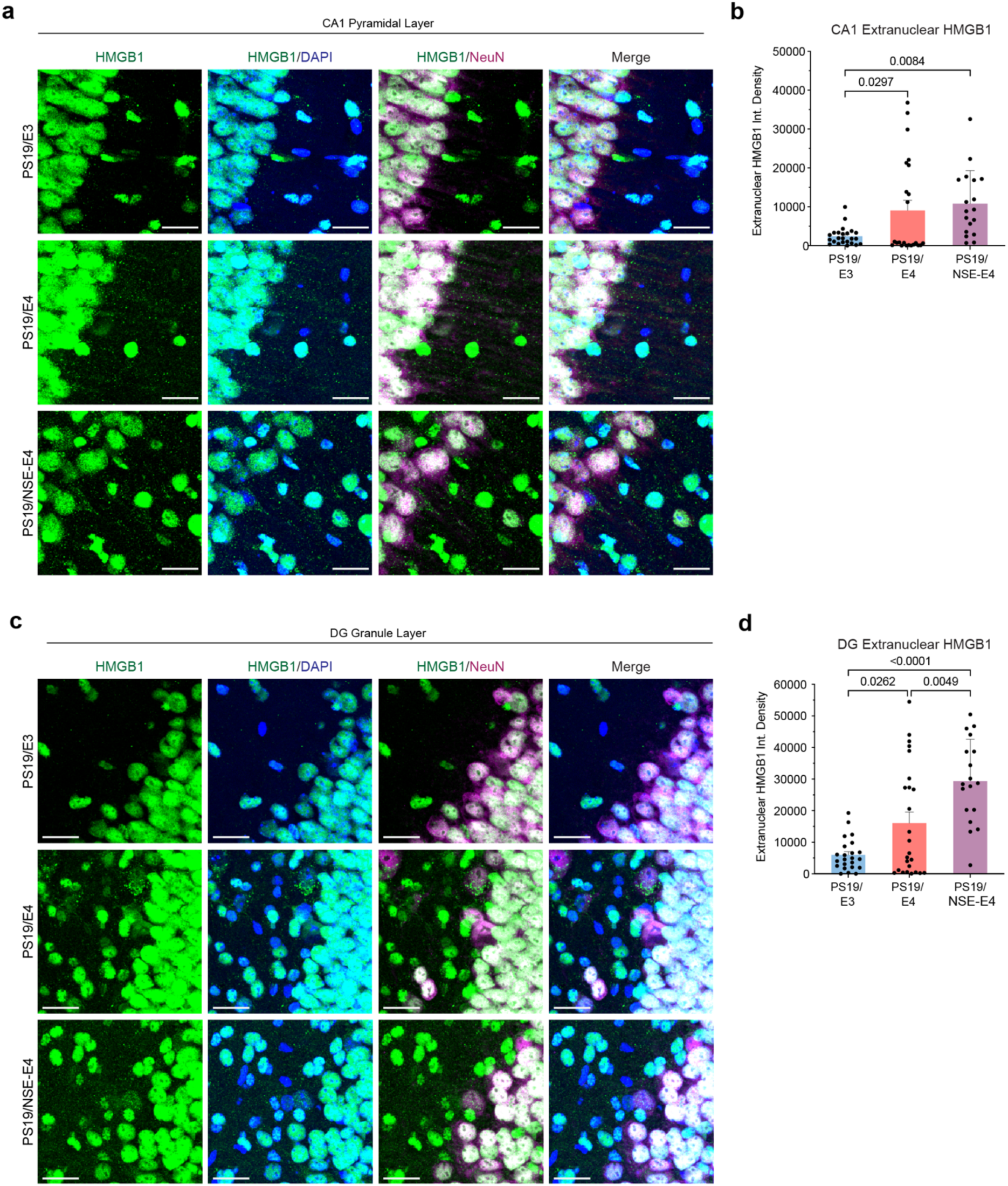
Neuronal APOE4 alone stimulates intracellular HMGB1 translocation. **a**, Representative images of CA1 pyramidal layer from 10-month-old PS19/E3, PS19/E4, and PS19/NSE-E4 mice, immunolabeled for HMGB1 (green), DAPI (blue), and NeuN (magenta) (scale bar, 20 µm). **b**, Quantifications of extranuclear HMGB1 fluorescent intensity in the CA1 of PS19/E3, PS19/E4, and PS19/NSE-E4 mice. **c**, Representative images of DG granule layer from PS19/E3, PS19/E4, and PS19/NSE-E4 mice, immunolabeled for HMGB1 (green), DAPI (blue), and NeuN (magenta) (scale bar, 20 µm). **b**, Quantifications of extranuclear HMGB1 fluorescent intensity in the DG of PS19/E3, PS19/E4, and PS19/NSE-E4 mice. In **b**,**d**, PS19/E3 *n* = 24; PS19/E4 *n* = 25; and PS19/NSE-E4 *n* = 18; data is expressed as mean ± s.e.m and was assessed via one-way analysis of variance (ANOVA) with Tukey’s post hoc multiple comparisons test.

### Neuronal APOE4 alone is sufficient to potentiate tau propagation and related pathologies

PS19 mice constitutively express mutant human tau, resulting in severe p-tau accumulation. To better determine how neuronal APOE4 influences initial tau seeding and propagation, we unilaterally injected adeno-associated virus-2 encoding P301S mutant tau (AAV2-tau-P301S) into the hippocampus of 10.5-month-old NSE-E4 mice (Fig. 5a). Knock-in mice carrying human APOE3 or APOE4 (referred to as APOE3 and APOE4 mice, respectively) were used as controls. To best observe tau spread, all mice contained only the wildtype mouse *Mapt* gene.

**Fig. 5.**
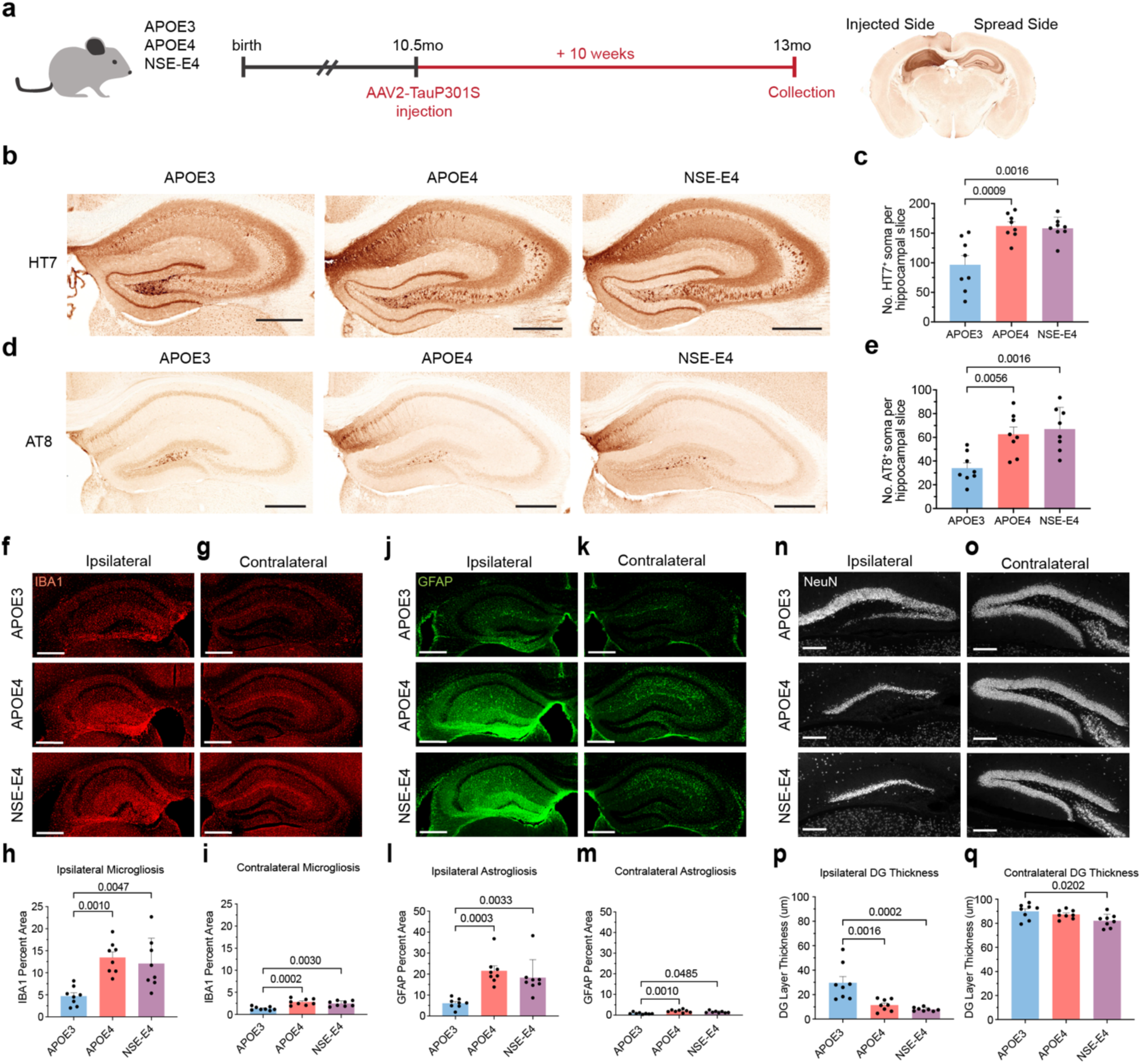
Neuronal APOE4 alone promotes Tau propagation and subsequent pathologies. **a**, Schematic depicting experimental design of *in vivo* Tau propagation study. **b**, Representative images of human Tau (HT7 monoclonal antibody) immunolabeling of the contralateral hippocampus of APOE3, APOE4, and NSE-E4 mice, 10 weeks post unilateral injection of mutant human Tau virus (scale bar, 500 µm). **c**, Quantifications of the number of HT7-positive soma in the contralateral hippocampus of these mice. **d**, Representative images of p-Tau (AT8) immunolabeling of the contralateral hippocampus of injected APOE3, APOE4, and NSE-E4 mice (scale bar, 500 µm). **e**, Quantifications of the number of AT8-positive soma in the contralateral hippocampus of these mice. **f-g**, Representative images of microglial (IBA1) immunolabeling of the ipsilateral (**f**) and contralateral (**g**) hippocampus of injected APOE3, APOE4, and NSE-E4 mice (scale bar, 500 µm). **h,i**, Quantifications of the percent area coverage of microglia (IBA1) in the ipsilateral (**h**) and contralateral (**i**) hippocampus of these mice. **j,k**, Representative images of astrocytic (GFAP) immunolabeling of the ipsilateral (**j**) and contralateral (**k**) hippocampus of injected APOE3, APOE4, and NSE-E4 mice (scale bar, 500 µm). **l,m**, Quantifications of the percent area coverage of astrocyte (GFAP) in the contralateral (**l**) and ipsilateral (**m**) hippocampus of these mice. **n,o**, Representative images of DG neuronal layers (NeuN) of the ipsilateral (**n**) and contralateral (**o**) hippocampus of injected APOE3, APOE4, and NSE-E4 mice (scale bar, 200 µm). **p,q**, Quantifications of the DG granule layer thickness in the contralateral (**p**) and ipsilateral (**q**) hippocampus of these mice. For quantifications in **c**,**e**,**h**,**i**,**l**,**m**,**p**,**q**, APOE3 *n* = 8; APOE4 *n* = 8; and NSE-E4 *n* = 8; data is expressed as mean ± s.e.m and was assessed via one-way analysis of variance (ANOVA) with Tukey’s post hoc multiple comparisons test.

Consistent with previous studies demonstrating the spread of intracranially-injected tau to anatomically connected brain regions^14,49–53^, all mice exhibited human tau- and p-tau-positive neuronal soma in the hippocampus contralateral to the injection site 10 weeks post-injection, confirming tau propagation (Fig. 5b-e). Interestingly, when compared with APOE3 controls, both APOE4 and NSE-E4 mice had over 60% more neurons positive for human tau (HT7), and more than 80% more neurons positive for p-tau (AT8) in the contralateral hippocampus (Fig. 5c,e). To ensure equivalent viral expression, we quantified human tau levels in the ipsilateral hippocampus, where HT7 signal proved to be comparable across all genotypes (Extended Data Fig. 3a,b). After normalizing contralateral counts to ipsilateral HT7 area coverage, AT8-positive and HT7-positive soma proportions remained significantly greater in the APOE4 and NSE-E4 groups (Extended Data Fig. 3c-f). Notably, pathological p-tau (AT8) was also elevated in the ipsilateral hippocampus of APOE4 and NSE-E4 mice relative to APOE3 mice (Extended Data Fig. 3d,e), consistent with the increased p-tau accumulation observed in our PS19/E4 and PS19/NSE-E4 tauopathy mice discussed above (Fig. 1a,b).

Because the entorhinal cortex (EC) is highly interconnected with the hippocampus and a known locus of trans-synaptic tau spread^39,52,53^, we also examined tau distribution in both ipsilateral and contralateral EC caudal to the injection site. Strikingly, NSE-E4 mice exhibited higher levels of both total human tau and p-tau in the ipsilateral EC relative to APOE3 mice (Extended Data Fig. 3g-i). Although tau-positivity was lower overall on the contralateral side given its extended distance from the original injection site, NSE-E4 mice also demonstrated higher total tau levels in contralateral EC compared to APOE3 controls (Extended Data Fig. 3j,k). Collectively, this suggests that neuronal APOE4 directly promotes tau spread, even across multiple nodes from the original injection site.

We next asked how neuronal APOE4-promoted tau spread influenced other AD pathologies. Immunostaining for microglia (IBA1) and astrocytes (GFAP) revealed significantly elevated gliosis in both ipsilateral and contralateral hippocampi of APOE4 and NSE-E4 mice compared with APOE3 mice (Fig. 5f-m). As expected, gliosis was most pronounced on the ipsilateral side (Fig. 5f-m), likely reflecting local inflammatory responses to high tau load, though lower-level gliosis was also visible in the contralateral side. Thus, within conditions of high tau levels, as exhibited by the ipsilateral side, or conditions of early tau propagation, as characterized on the contralateral side, neuronal APOE4 alone proved sufficient to induce gliosis.

We then investigated the effect of neuronal APOE4-promoted tau spread on neurodegeneration by quantifying neuronal layer width and hippocampal volume. On the ipsilateral side, where tau expression was highest, we found severe DG granule cell loss, particularly of the ventral DG blade (Fig. 5n,o). Despite extensive degeneration across all genotypes, APOE4 and NSE-E4 mice exhibited greater neuronal loss relative to APOE3 mice (Fig. 5n,o). Moreover, ipsilateral hippocampal volume was also significantly reduced in the APOE4 and NSE-E4 mice (Extended Data Fig. 3l-n). Even on the contralateral side, NSE-E4 mice displayed a modest but significant reduction in neuronal layer thickness relative to APOE3 mice (Fig. 5p,q), indicating that APOE4, especially neuronal APOE4, potentiates degeneration downstream of tau pathology and tau-associated gliosis. Together, these findings demonstrate that neuronal APOE4 alone is sufficient to drive initial tau propagation, neuroinflammation, and subsequent neurodegeneration across interconnected brain regions.

### Neuronal APOE4 alone is sufficient to prompt loss of vulnerable neuronal subtypes

Given the depth and resolution offered by modern transcriptomics profiling, we next performed single-nucleus RNA sequencing (snRNA-seq) on hippocampi of 10-month-old PS19/E3, PS19/E4, and PS19/NSE-E4 mice to investigate cell type-specific changes due to neuronal APOE4 expression. After quality control and normalization, we obtained 98,610 nuclei expressing a collective 26,195 unique transcripts. With clustering using Louvain algorithms and visualization with uniform manifold approximation and projection (UMAP), we identified 33 unique cell clusters (Fig. 6a). Based on canonical marker gene expression (Extended Data Fig. 4a,b), we identified eleven excitatory neuron clusters (3, 4, 7, 10, 16, 18, 27, 28, 29, 31, 32), three inhibitory neuron clusters (5, 11, 30), seven subiculum neuron clusters (6, 9, 12, 19, 20, 22, 24), and twelve non-neuronal clusters, including three oligodendrocyte clusters (1, 2, 17), two microglia clusters (8, 13), two oligodendrocyte progenitor cell (OPC) clusters (14, 33), two astrocyte clusters (15, 23), and one choroid plexus cluster (26) (Fig. 6a, Extended Data Fig. 4a,b and Supplementary Table 1).

**Fig. 6.**
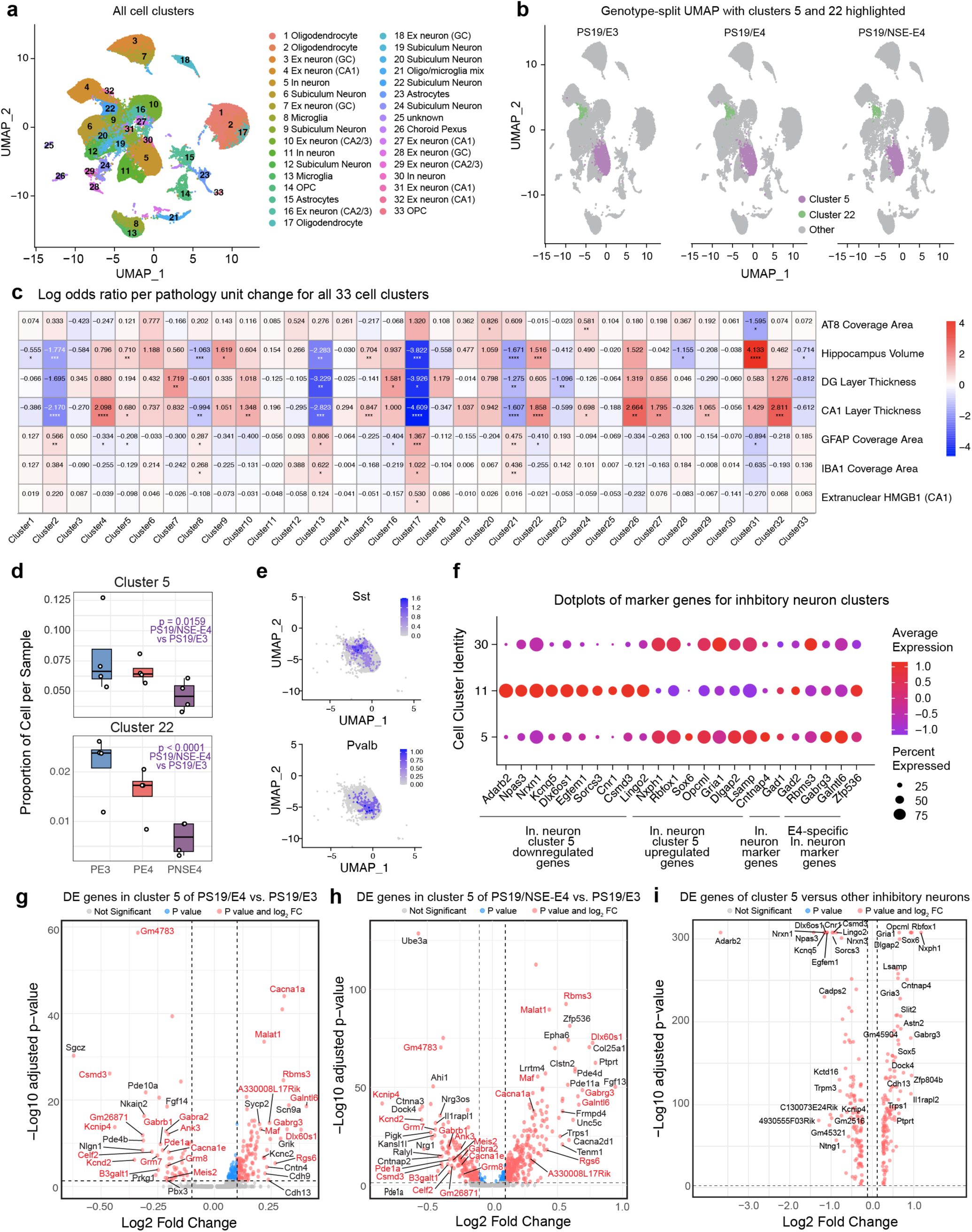
Neuronal APOE4 alone elicits loss of vulnerable neuronal subtypes. **a**, UMAP plot of the 33 unique cell clusters identified in hippocampi from 10-month-old PS19/E3 (*n* = 4), PS19/E4 (*n* = 4), and PS19/NSE-E4 (*n* = 4) mice. **b**, Genotype-split UMAP highlighting neuronal clusters 5 and 22 across each genotype group. **c,** Heatmap plot of the log odds ratio (LOR) per unit change for each pathological parameter against each cell cluster. The LOR represents how changes in each pathological measurement (AT8 coverage area, hippocampal volume, etc.) are associated with the odds that a cell belongs to a given cluster versus not, while accounting for genotype and sample variability through random effects. Associations with pathologies are on a colored scale (negative associations, blue; positive associations, red). **d**, Box plot of the proportion of cells from each sample for clusters 5 and 22 (PS19/E3, *n* = 4; PS19/E4, *n* = 4; and PS19/NSE-E4, *n* = 4). From bottom to top, the hinges of the box plots correspond to the 25^th^, 50^th^, and 75^th^ percentiles. The upper and lower whiskers of the box plot extend to the largest and smallest values, respectively, though no further than 1.5 x IQR from the nearest hinge. IQR, interquartile range, or distance between the 25^th^ and 75^th^ percentiles. The LOR are the mean ± s.e.m estimates for these clusters, which represents the change in the log odds of cells per sample from each mouse belonging to the respective clusters relative to log odds of cells per sample from PS19/E3 mice. **e**, Feature plots illustrating the expression of SST and PV in inhibitory neuron cluster 5. **f**, Dot-plot of normalized average expression of marker genes and genes of interest for selected inhibitory neuron clusters. The size of the dots is proportional to the percentage of cells expressing a given gene. Average expression is on a colored scale (lower expression, blue; higher expression, red). **g,** Volcano plot of the differentially-expressed (DE) genes between PS19/E4 and PS19/E3 in cluster 5 inhibitory neurons. **h**, Volcano plot of the DE genes between PS19/NSE-E4 and PS19/E3 in cluster 5 inhibitory neurons. Up- or down-regulated genes shared between **g** and **h** are indicated in red font. **i**, Volcano plot of the DE genes between inhibitory neuron cluster 5 and all other inhibitory neuron clusters. In **g-i**, Dashed lines represent log_2_ fold change threshold of 0.1 and an adjusted p value threshold of 0.05. The unadjusted P values and log_2_ fold change values used were generated from the differential expression analysis using the two-sided Wilcoxon rank-sum test as implemented in the FindMarkers function of the Seurat package. In **c,** unadjusted p values are from fits to a GLMM_histopathology. In **d**, unadjusted p values are from fits to a GLMM_AM. Two-sided association tests were used. Ex neuron, excitatory neuron; In neuron, inhibitory neuron; oligo, oligodendrocyte.

To identify the clusters most altered by disease, we applied a generalized linear mixed-effects model (GLMM) to determine the log odds ratio (LOR) between each cluster and the neuropathological measures reported above, including AT8 coverage, hippocampal volume, DG and CA1 neuronal layer thickness, GFAP and IBA1 coverage, and extranuclear HMGB1 levels. Several neuronal clusters were negatively associated with pathology, including inhibitory cluster 5 and subiculum cluster 22, both of which correlated positively with hippocampal volume and CA1 layer thickness but negatively with astrocytic GFAP levels (Fig. 6b,c and Supplementary Table 2). Notably, LOR analysis of the cell proportions revealed that PS19/NSE-E4 mice had a significantly reduced proportion of cells belonging to neuron clusters 5 and 22 relative to the PS19/E3 controls (Fig. 6b,d and Supplementary Table 2), suggesting selective loss of these neuronal populations in the PS19/NSE-E4 model. These findings indicate that neuronal APOE4 alone is sufficient to drive vulnerability and degeneration of specific neuronal subtypes. Previous reports have highlighted somatostatin-positive (SST) neurons as particularly susceptible in AD, with SST neuron loss being highly associated with AD and APOE4 expression^54–57^. Consistent with this, cluster 5 contained a high proportion of SST-expressing neurons (Fig. 6e), further confirming it as a neuronal APOE4-induced disease-vulnerable inhibitory neuron class.

We next examined transcriptional changes within these vulnerable clusters (Fig. 6f-i, Extended Data Fig. 5a-d and Supplementary Table 3). Differential expression analysis of cluster 5 revealed that, relative to PS19/E3 mice, both PS19/NSE-E4 and PS19/E4 mice showed reduced expression of ion channel genes regulating excitability, including *Kcnd2*^58^ and *Kcnip4*^59^, and synaptic organization genes, such as *Nrgn* and *Grm7* (Fig. 6g,h and Supplementary Table 3). When compared to other inhibitory neuron clusters, cluster 5 demonstrated downregulation of additional synaptic and excitability genes, including *Nrxn1*, *Sorcs3*, and *Kcnq5* (Fig. 6f,i and Supplementary Table 3), suggesting that impaired inhibitory signaling and synaptic maintenance may underlie its vulnerability. Furthermore, cluster 5 upregulated *Cntnap4*, which encodes a synaptic protein involved in inhibitory signal transmission and whose deficiencies have been linked to cognitive impairments and p-tau accumulation^60,61^. Thus, neuronal APOE4 expression appears sufficient to disrupt inhibitory circuit integrity, predisposing the hippocampus to excitotoxicity.

When compared to other subiculum clusters, subiculum cluster 22 also displayed similar changes, including downregulation of *Dpp10*, *Lrrc4c*, and *Nrg1*—genes involved in potassium channel regulation and synaptic organization—and upregulation of *Hcn1*, *Gria1*, and *Grm5*, indicative of hyperexcitability (Extended Data Fig. 5a–d and Supplementary Table 3). Cluster 22 also showed reduced *PTPRD* expression (Extended Data Fig. 5a,d and Supplementary Table 3), mutations of which have been previously linked to neurofibrillary tangle (NFT) accumulation^62^. Given that the subiculum bridges the EC (where NFTs first emerge in AD) with the hippocampus, these neurons may be playing a key role in trans-synaptic tau propagation. Interestingly, *Cntnap5a* and *Cntnap5b* were upregulated in this cluster (Extended Data Fig. 5a,d and Supplementary Table 3), with further upregulation in PS19/NSE-E4 and PS19/E4 mice relative to PS19/E3 controls (Extended Data Fig. 5b,c and Supplementary Table 3). As the *Cntnap5* family is predicted to be involved in proteolysis^63^, this may represent a compensatory response to tau accumulation. Taken together, these data demonstrate that neuronal APOE4 alone is sufficient to drive selective loss and dysfunction of vulnerable neuronal populations.

### Neuronal APOE4 alone is sufficient to potentiate disease-associated glial phenotypes

Given that neuronal APOE4 alone was able to drive gliosis (Fig. 3), we next examined the transcriptomic signatures of glial populations. In contrast to the neuronal clusters described above, microglia clusters 8 and 13 and oligodendrocyte clusters 2 and 17 appeared disease-associating, correlating positively with gliosis and negatively with hippocampal volume (Fig. 6c and Supplementary Table 2). Cell proportion analysis revealed that the proportion of cells in microglia cluster 8 and oligodendrocyte cluster 17 were enriched in PS19/NSE-E4 mice relative to PS19/E3 controls, with microglia cluster 13 displaying a similar trend (Fig 7a-c and Supplementary Table 2). These results suggest that APOE4, even when restricted to neurons, promotes enrichment of disease-associated microglia and oligodendrocyte populations. Interestingly, oligodendrocyte cluster 17 also correlated positively with extranuclear localization of HMGB1 in the CA1 region (Fig. 6c and Supplementary Table 2), suggesting that these disease-associated oligodendrocytes may be directly responsive to neuronal APOE4-induced release of proinflammatory HMGB1^18^.

**Fig. 7.**
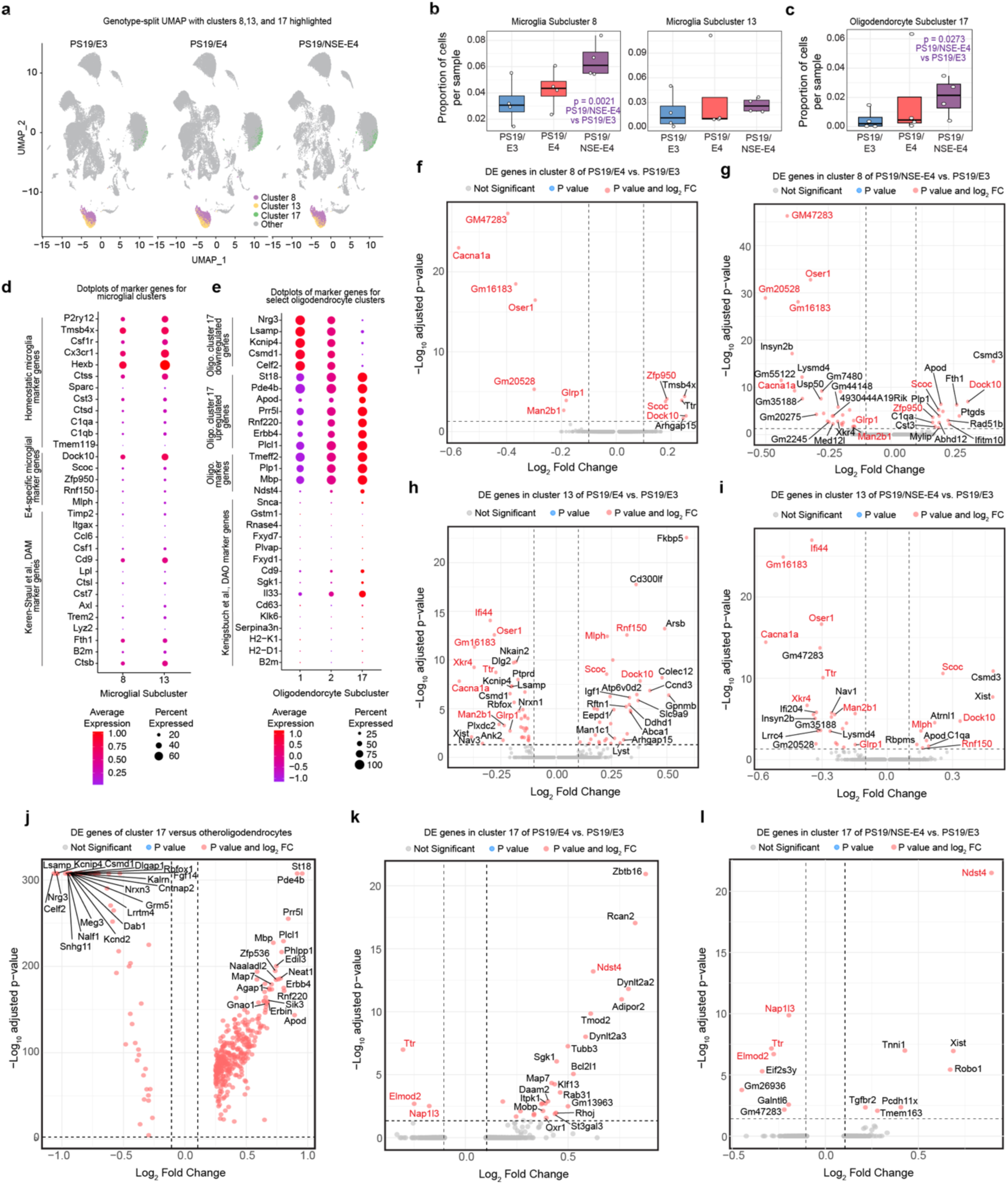
Neuronal APOE4 alone produces disease-associated microglia and oligodendrocytes. **a**, Genotype-split UMAP highlighting microglial clusters 8 and 13 and oligodendrocyte cluster 17 across each genotype group. **b,c**, Box plot of the proportion of cells from each sample for microglial clusters 8 and 13 (**b**) and oligodendrocyte cluster 17 (**c**) (PS19/E3, *n* = 4; PS19/E4, *n* = 4; and PS19/NSE-E4, *n* = 4). From bottom to top, the hinges of the box plots correspond to the 25^th^, 50^th^, and 75^th^ percentiles. The upper and lower whiskers of the box plot extend to the largest and smallest values, respectively, though no further than 1.5 x IQR from the nearest hinge. IQR, interquartile range, or distance between the 25^th^ and 75^th^ percentiles. The log odds ratios (LOR) are the mean ± s.e.m estimates of LOR for these clusters, which represents the change in the log odds of cells per sample from each mouse belonging to the respective clusters relative to log odds of cells per sample from PS19/E3 mice. **d,** Dot-plot of average expression of marker genes and genes of interest for microglial clusters. To best compare only two clusters, dot-plot values are presented without normalization. **e**, Dot-plot of normalized average expression of marker genes and genes of interest for oligodendrocyte clusters (**e**). For **d**,**e**, the size of the dots is proportional to the percentage of cells expressing a given gene. Average expression is on a colored scale (lower expression, blue; higher expression, red). **f**, Volcano plot of the differentially-expressed (DE) genes between PS19/E4 and PS19/E3 in cluster 8 microglia. **g**, Volcano plot of the DE genes between PS19/NSE-E4 and PS19/E3 in cluster 8 microglia. Up- or down-regulated genes shared between **f** and **g** are indicated in red font. **h**, Volcano plot of the differentially-expressed (DE) genes between PS19/E4 and PS19/E3 in cluster 13 microglia. **i**, Volcano plot of the DE genes between PS19/NSE-E4 and PS19/E3 in cluster 13 microglia. Up- or down-regulated genes shared between **h** and **i** are indicated in red font. **j**, Volcano plot of the DE genes between oligodendrocyte cluster 5 and all other oligodendrocyte clusters. **k**, Volcano plot of the DE genes between PS19/E4 and PS19/E3 in cluster 17 oligodendrocytes. **l**, Volcano plot of the DE genes between PS19/NSE-E4 and PS19/E3 in cluster 17 oligodendrocytes. Up- or down-regulated genes shared between **k** and **l** are indicated in red font. In **f**-**l**, dashed lines represent log_2_ fold change threshold of 0.1 and adjusted p value threshold of 0.05. The adjusted p values and log_2_ fold change values used were generated from the differential expression analysis using the two-sided Wilcoxon rank-sum test as implemented in the FindMarkers function of the Seurat package. Unadjusted p values in **b**,**c** are from fits to a GLMM_AM; association tests were two-sided.

We next characterized the transcriptional changes occurring in these disease-associating glial populations (Fig. 7d,e and Supplementary Table 3). Both microglial clusters 8 and 13 expressed canonical disease-associated microglia (DAM) markers, including *Cd9*, *Fth1*, and *Ctsb* (Fig. 7d), though expression levels were highest in cluster 13. Cluster 13 also upregulated additional DAM genes, including *Csf1*, *Lpl*, *Ctsl*, *Axl*, and *B2m* (Fig. 7d). Relative to PS19/E3 microglia, both PS19/E4 and PS19/NSE-E4 microglia upregulated *Dock10, Scoc, Zfp950*, and *Rsnf150*, and *Mlph*, while down-regulating *Cacna1a, Ifi44, Man2b1, and Xkr4* (Fig. 7f-i and Supplementary Table 3), suggesting broad shifts in microglia activation and function. Interestingly, both microglia clusters 8 and 13 also contained substantial proportions of cells expressing homeostatic markers, including *P2ry12*, *Csf1r*, *Cx3cr1*, and *Hexb*. Given the projected role of microglial APOE in DAM activation^40,64^, these APOE-deficient microglia may reside in a novel or partially activated state.

Oligodendrocyte cluster 17 also exhibited disease-associated transcripts, with increased expression of traditional disease-associated-oligodendrocyte (DAO) genes, such as *1l33*, *Sgk1*, and *Cd9* (Fig. 7e). Relative to the other oligodendrocyte clusters, cluster 17 upregulated multiple genes, including *St18*, *Pde4b*, *Apod*, *Prr5l*, *Rnf220*, *Erbb4*, and *Ndst4* (Fig. 7e,j and Supplementary Table 3). Interestingly, several of these upregulated genes are implicated in AD, as *St18* has previously been found to associate with cognitive decline in AD^65^, *Erbb4* is elevated in AD brains^66^, and *Pde4b* inhibition improves cognitive function in APP mouse models^67,68^. Meanwhile, genes involved in axonal support, neuronal communication, and immune regulation, including *Nrg3*, *Lsamp*, *Kcnip4*, *Csmd1*, and *Celf2,* were downregulated (Fig. 7e,j-l and Supplementary Table 3). Intriguingly, *Nrg3* mutations have been previously linked to AD^69^. PS19/NSE-E4 and PS19/E4 mice also further downregulated *Elmod2*, *Nap1l3*, and *Ttr* (Fig. 7j-k and Supplementary Table 3), the latter of which is found in lower levels in AD patient samples^70–73^ and thought to play a neuroprotective role in AD^74,75^. Given these altered gene expression patterns, neuronal APOE4 alone proved sufficient to drive disease-associated oligodendrocyte signatures.

We also examined the two astrocyte clusters 15 and 23 (Extended Data Fig. 6a-g and Supplementary Table 2). Cluster 15 appeared more health-associating and homeostatic, as it correlated positively with hippocampal volumes and CA1 neuronal layer thickness (Fig. 6c and Supplementary Table 2), and made up a higher proportion of cells in PS19/E3 relative to PS19/NSE-E4 mice (Extended Data Fig. 6b and Supplementary Table 2). Consistent with this, PS19/E4 and PS19/NSE-E4 cluster 15 astrocytes upregulated *Abca1* (Extended Data Fig. 6c,d and Supplementary Table 3), loss-of-function mutations of which are largely associated with AD. In contrast, cluster 23 demonstrated some association with pathology, correlating negatively with DG layer thickness (Fig. 6c and Supplementary Table 2). While cluster 23 astrocytes displayed similar cell proportions and limited differential expression across genotypes (Extended Data Fig. 6e,f and Supplementary Table 3), ApoD – a protein known to be elevated in AD brains^76^ - was upregulated in both PS19/E4 and PS19/NSE-E4 hippocampi relative to PS19/E3 controls. Notably, both astrocyte clusters exhibited minimal expression of disease-associated astrocyte (DAA) markers, but cluster 15 retained higher expression of homeostatic genes such as *Luzp2*, *Slc1a2*, *Gfap*, *Aqp4*, *Gja1*, and *Slc7a10* relative to cluster 23 (Extended Data Fig. 6c), suggesting cluster 15 represents homeostatic astrocytes. Collectively, these results demonstrate that in the absence of glial APOE, neuronal APOE4 expression can drive transcriptional activation of microglial and oligodendritic disease states, while subtly reshaping astrocyte populations toward both activated and homeostatic subtypes.

## Discussion

In this study, we investigated whether neuron-specific expression of APOE4 is sufficient to drive AD pathogenesis. We found that selective neuronal APOE4 expression recapitulates, and in some cases exceeds, the pathologies observed in models with global APOE4 expression. The PS19/NSE-E4 model exhibited p-tau accumulation and progression, hippocampal atrophy, neuron loss, gliosis, HMGB1 translocation, and transcriptomic disease signature changes comparable to the PS19/E4 model. These findings establish that neuronal APOE4 alone is sufficient to initiate and exacerbate the key pathological hallmarks of AD.

Interestingly, we observed multiple phenotypes that were more severe in the PS19/NSE-E4 and NSE-E4 mice relative to the PS19/E4 and APOE4 lines, respectively, including CA1 degeneration (Fig. 2d,f), astrogliosis (Fig. 3e-h), HMGB1 translocation in the DG (Fig. 4c-d), and tau spread into the EC (Extended Data Fig. 3g-k). Given that neurons secrete only about 10% of the APOE they produce^77^, much of the APOE4 produced in the PS19/NSE-E4 model will accumulate intraneuronally, where it could overwhelm the proteosome or directly influence HMGB1 translocation or tau phosphorylation. Additionally, neuronal APOE4 is known to produce neurotoxic fragments^13,30,78^, which could further potentiate tau phosphorylation, neurodsyfunction, and subsequent pathologies.

Viewing this study in conjunction with previous work showing that selectively removing neuronal APOE4 prevents the primary AD pathological hallmarks^14^, our data indicate that neuronal APOE4 is both necessary and sufficient for AD pathogenesis. Nevertheless, other cellular sources of APOE4 could still contribute to disease severity. As mentioned previously, astrocytes are the greatest producers of APOE in the brain; however, other CNS cells, including neurons, microglia, and vascular cells, also express APOE^5^. Other cell type-specific APOE deletion studies have found some protective effects when eliminating astrocytic APOE, including reduced Aβ accumulation in an APP/PS1-21 model^79^ and attenuated degeneration, glial disease signatures, and tau pathology in a PS19 model^80^. Similarly, ablation of microglial APOE in the APP/PS1 model also diminishes Aβ plaques^81^. Additionally, vascular APOE4 expression has been linked to impaired blood flow and spatial learning^82^. Collectively, these studies suggest other cellular sources of APOE4 may exacerbate pathologies downstream, likely through amplifying glial responses and downstream neurodegeneration, as outlined previously in the Cell Type-Specific APOE4 Cascade Model of AD^5^. Importantly, our PS19/NSE-E4 model, which exhibits key AD hallmarks without any non-neuronal APOE4, demonstrates that neuronal APOE4 alone is sufficient to initiate and drive AD pathologies.

Despite lacking microglial APOE expression, PS19/NSE-E4 mice exhibited increased levels of both general and phagocytic microglia relative to the PS19/E3 mice (Fig. 3a-d). Several traditional DAM genes, including *Cd9*, *Fth1*, and *Ctsb*, also appear elevated in microglia clusters 8 and 13, which are highly associated with the PS19/NSE-E4 group and neurodegenerative pathologies (Fig. 7b,d). These findings align with a previous study that found microglial abundance and transcriptional signatures were largely preserved between a 5xFAD model with microglia-specific APOE-KO and the control 5xFAD mice^83^. Microglial APOE may therefore not be necessary for transcriptional shifts into a DAM state. Alternatively, given the high expression of homeostatic markers also in clusters 8 and 13, as well as recent research highlighting how microglia reactivity may function more as a spectrum than a binary active or resting state^84^, these APOE-deficient microglia may instead represent a partially activated or intermediate state.

The demonstration that neuronal APOE4 alone can trigger gliosis, neurodegeneration, and tau pathology implies that targeting neuronal APOE4 or its downstream signaling pathways would be a more precise therapeutic strategy than global APOE4 modulation, which could cause unanticipated side effects. Current anti-APOE4 immunotherapies^3,85–89^ could be adapted for neuron-specific delivery using viral vectors^90,91^. Viral vectors targeting neurons could also be used in conjunction with gene editing techniques. For example, because APOE2 greatly reduces AD risk compared with APOE4 and APOE3^92,93^, and APOE2 gene therapy in mice reduces amyloid plaques and synaptic loss^94^, converting neuronal APOE4 to APOE2 using CRISPR technologies could provide protection. Similarly, the R136S (Christchurch) mutation, even when edited onto an APOE4 background, has proven protective in PS19/E4 mice^15^, and thus could offer another promising therapeutic avenue. Since neuron-specific elimination of APOE in PS19 mice is sufficient to ameliorate tau pathology, neurodegeneration, and neuroinflammation^14^, CRISPR-interference (CRISPRi) strategies would also prove effective. Beyond viral delivery, antisense oligonucleotides (ASO) against APOE^95,96^ could be adapted for neuronal delivery via ligand-conjugation. Alternatively, ASOs targeting APOE-intron 3 (APOE-I3), an APOE splicing variant found exclusively in neurons^97^, may offer an even more efficient therapy against neuronal APOE4.

In addition to directly lowering neuronal APOE4 levels, interventions targeting downstream consequences of neuronal APOE4 would also be promising. As demonstrated in our PS19/NSE-E4 analysis and in other studies, neuronal APOE4 drives inhibitory neuron vulnerability^98^. As inhibitory neuron loss contributes to hyperexcitability^99–101^, human iPSC-based stem-cell replacement therapies could be used to combat this neurodysfunction^102^. Furthermore, as neuronal APOE4 produces more neurotoxic fragments^13^, reducing fragments by developing protease inhibitors capable of counteracting APOE4 cleavage in neurons may help reduce neuronal-APOE4-induced pathologies. Ultimately, multiple therapeutic modalities could be employed to ameliorate the effects of neuronal APOE4.

In summary, our findings highlight neuronal APOE4 as a central initiator of AD-related pathologies, driving p-tau accumulation and propagation, hippocampal and neuronal loss, neuroinflammation, and disease-signature changes across neuronal and glial cell types. These results position neuronal APOE4 and its immediate downstream effects as central targets for therapeutic intervention in APOE4-related AD.

## METHODS

### Mice

NSE-APOE4 (NSE-E4) mice were generated previously on an APOE-KO background to ensure exclusive APOE expression by neurons^28,29,33^. The NSE-APOE4 mice without mouse ApoE were crossbred with PS19 transgenic mice (B6;C3-Tg(Prnp-MAPT*P301S)PS19Vle/J) (The Jackson Laboratory, 008169) to generate PS19/NSE-E4 mice solely expressing APOE4 in neurons. PS19/E3 and PS19/E4 mice generated previously in-house^14,15^ were created via crossing human APOE4 (E4) and APOE3 (E3) knock-in mice^103^ with PS19 transgenic mice^36^. These APOE knock-in mice contain a LoxP-floxed human APOE gene for use in other studies. All mice were produced on a C57BL/6 genetic background. Mice were housed in a pathogen-free barrier facility with a 12 hr light cycle at 19-23°C and 30-70% humidity. Animals were identified by ear punch and genotyped by polymerase chain reaction of ear or tail clippings. All animals received only procedures reported in this study. For all studies, male and female mice were used. All animal experiments were conducted in accordance with the guidelines and regulations of the National Institute of Health, the University of California, and the Gladstone Institutes under the protocol AN205544. All protocols and procedures followed the guidelines of the Laboratory Animal Resource Center and the University of California, San Francisco (UCSF) and the ethical approval of the UCSF institutional animal care and use committee.

### Stereotaxic mouse surgery

NSE-E4, E4, and E3 mice underwent intracranial injection of tau virus into the hippocampus for *in vivo* tau spread study. At approximately 10.5 months of age, mice were anesthetized and secured in a stereotaxic frame with a nose cone (Kopf instruments Model 940). Scalp was shaven and sterilized before a midline incision was made. After localizing Bregma, a stereotaxic site was drilled with a 0.5 mm microburr (Fine Science Tools, Cat. #19007-05) using coordinates X = +1.5 and Y = −2.1. Injection needle was slowly lowered to Z = -2.1 (to target the DG region), and 2 μl of Adeno-associated virus-2 (AAV2) expressing pathological human tau-P301S (AAV2(Y444F)-smCBAhuman_P301S_tau-WPRE; Virovek) concentrated at 2.10E + 13 vg/ml was injected at 0.5 ul/min. After allowing the virus to diffuse for 3 minutes following injection, needle was removed and scalp was sutured. Local anesthetic (Lidocaine; Vedco) was applied subcutaneously along the incision, and buprenorphine (Henry Schein, Cat. #55175) and ketofen (Henry Schein, Cat. #005487) were administered for analgesia. Mice were monitored on a heating pad until ambulatory and then single-housed for the duration of the study.

### Mouse tissue collection

For characterization of the PS19/NSE-E4 line, mouse brains were collected at 10 months of age as previously described^14,15^. In brief, mice were deeply anesthetized by intraperitoneal injection of avertin (Sigma-Aldrich, Cat. #T48402) and transcardially perfused with 0.9% saline for 2 minutes. Brains were dissected out of the skull and bisected into two hemispheres. Hippocampus was dissected out from the left hemisphere and snap frozen on dry ice before storing at -80°C. Right brain hemisphere was drop-fixed in 4% PFA (Electron Miscropscopy Science, Cat. #15710-S) for 48 hours, washed in PBS (Thermo Scientific, Cat. #J75889K2), and placed in 30% sucrose (Sigma-Aldrich, Cat. #S7903) for 48 hours to prepare for sectioning. Fixed hemi-brains were sliced into 30 µm coronal sections using a sliding microtome (Leica, Cat. #SM2010R). Sections were stored in cryoprotectant solution (30% ethylene glycol, Fisher Chemical, Cat. #E178-4; 30% glycerol, Sigma-Aldrich, Cat. #G9012-1GA; and 40% 1x PBS) at -20°C. For the tau spread study, mouse brains were collected at 13 months of age as described above, with the whole brain drop-fixed to examine contralateral tau spread. A dorsal notch was added to the left hemisphere during sectioning to distinguish left-right orientation. Sections were stored in cryoprotectant solution at - 20°C.

### Immunohistochemistry

For immunofluorescence staining, several 30 μm-thick brain sections were washed three times with PBS-T (PBS + 0.1% Tween-20; Millipore Sigma, Cat. #P2287) and incubated in boiling Tris buffer (pH 8, Corning, Cat. #46-031-CM) for 15 minutes. Sections underwent three more PBS-T washes before blocking with 5% normal donkey serum (NDS; Jackson ImmunoResearch, Cat. #017000121) made in PBS-Tx (PBS + 0.2% TritonX; Millipore Sigma, Cat. #T8787) for 1 hour. Sections were then incubated in Mouse on Mouse (MOM; Vector Laboratories, Cat. #MKB-2213-1) blocking buffer (one drop MOM per 4ml of PBS-T) to block endogenous mouse staining and incubated overnight at 4°C with the following primary antibodies: anti-APOE (gt) 1:500 (Sigma-Aldrich, Cat. #178479); anti-APOE (rb) 1:500 (Cell Signaling, Cat. #13366); anti-CD68 1:200 (Bio-Rad, Cat. #MCA1957); anti-GFAP 1:750 (Sigma-Aldrich, Cat. #MAB3402); anti-HMGB1 1:200 (Abcam, Cat. #ab18256); anti-IBA1 (rb) 1:1000 (Abcam, Cat. #ab178846); anti-NeuN 1:500 (Sigma-Aldrich, Cat. #ABN90); anti-Olig2 1:100 (Abcam, Cat. #ab109186); and anti-S100β 1:200 (Abcam, Cat. #ab54642). All primary antibodies were diluted in 3% NDS made in PBS. The following day, sections were washed three times with PBS-T and incubated in the dark for 1 hour at room temperature with the following secondary antibodies: Donkey anti-goat 405 1:1000 (Invitrogen, Cat. #A48259); Donkey anti-goat 594 1:1000 (Invitrogen, Cat. #A11058); Donkey anti-guinea pig 647 1:1000 (Jackson Immuno, Cat. #706-605-148); Donkey anti-mouse 488 1:1000 (Invitrogen, Cat. #A-21202); Donkey anti-mouse 594 1:1000 (Invitrogen, Cat. #A-21203); Donkey anti-mouse 647 1:1000 (Invitrogen, Cat. #A-31571); Donkey anti-rabbit 488 1:1000 (Invitrogen, Cat. #A-21206); Donkey anti-rabbit 594 1:1000 (Invitrogen, Cat. #A-21207); Donkey anti-rat 488 1:1000 (Invitrogen, Cat. #A-21208); All secondary antibodies were diluted in 3% NDS made in PBS, with added DAPI (1:20,000; Thermo Scientific, Cat. #62248). After secondary incubation, sections were washed with PBS-T and mounted onto microscope slides (Fisher Scientific, Cat. #22-037-246). Slides were coverslipped with ProLong Gold mounting medium (Invitrogen, Cat. #P36930) and allowed to dry before imaging on an Aperio VERSA slide scanning microscope (Leica) at x10 or a FV3000 confocal laser scanning microscope (Olympus) at x20 or x60.

For diaminobenzidine (DAB) staining, several 30 um thick brain sections were washed three times with PBS-T incubated in boiling citrate buffer (10 mM sodium citrate; Fisher Bioreagents, Cat. # BP327-1) for 15 minutes, and washed thrice more with PBS-T before incubating in endogenous peroxidase block (3% H2O2; Sigma-Aldrich, Cat. #H1009-500mL, and 10% methanol; Fisher Scientific, Cat. #A412SK-4, in PBS) for 15 minutes. Sections underwent three more PBS-T washes before blocking with 10% NDS made in PBS-Tx for 1 hour. Sections were then incubated in avidin and biotin blocks (3 drops of each; Vector Laboratories, Cat. #SP-2001) for 15 minutes and MOM Blocking buffer (one drop MOM per 4ml of PBS-T) for 1 hour. Sections were incubated overnight at 4°C with the following primary antibodies: anti-AT8 1:100 (Invitrogen, Cat. #MN1020); anti-HT7 1:200 (Invitrogen, Cat. #MN1000). All primary antibodies were diluted in 3% NDS made in PBS-T.

The next day, sections were washed three times with PBS-T and incubated for 1 hour at room temperature with biotinylated donkey anti-mouse secondary (1:2000; Jackon ImmunoResearch, Cat. #715-065-150) diluted in 3% NDS in PBS-T. Sections were rinsed three times with PBS-T, incubated with ABC buffer (2 drops A and B per 5ml PBS; Vector Laboratories, Cat. #PK-6100) for 1 hour, and rinsed three times with PBS, before 2-minute incubation in DAB buffer (2 drops buffer stock solution, 4 drops DAB, and 2 drops H2O2 in 5ml milliQ H2O; Vector Laboratories, Cat. #SK-4100). Sections were immediately washed three times with milliQ H2O and then washed with PBS and mounted onto microscope slides. Slides were allowed to dry overnight and then submerged in two washes of xylene (Fisher Scientific, Cat. #HC7001GAL). Slides were coverslipped with DPX mounting medium (Sigma-Aldrich, Cat. #06522) and imaged with Aperio VERSA slide scanning microscope (Leica) at x10.

## Immunohistochemical quantification analysis

To exclude the possibility of bias for all quantifications, researchers were blinded to samples and automated image analysis programs were developed and used. For each analysis measuring immunostaining as a percentage of hippocampal area covered, several brain slices were stained and imaged according to the protocol above. After imaging, the hippocampal region was manually traced in Fiji (ImageJ) and a standardized threshold was used to determine the percent of positive staining area in the hippocampal region of interest.

To categorize p-tau staining types, two or three brain slices were stained with anti-AT8 following the DAB staining protocol described above, and categorization was performed manually, based on qualitative assessment of p-tau staining in the mossy fibers, the DG granule layer, CA3 pyramidal layer, and the stratum radiatum.

For counting tau-positive soma in *in vivo* tau spread cohort, two brain slices were stained with anti-AT8 or anti-HT7 in accordance with the DAB staining protocol described above. Hippocampal and EC regions were manually traced. For HT7 staining, subtract background was used to better distinguish cell soma. Count was then determined using standardized size and circularity parameters with the analyze particle function.

For DG granule cell layer and CA1 pyramidal layer thickness measurements, two brain sections were stained according to the protocol above with anti-NeuN to highlight the cell layers. The thickness was measured manually in Fiji as follows: a line was drawn perpendicular to the NeuN cell layer at 4 locations in both sections, and an average was taken for each mouse.

To measure nuclear and extra-nuclear levels of HMGB1, two brain slices were labeled with anti-HMGB1, anti-NeuN, and DAPI as described in the protocol above. Images were analyzed in Fiji, and a standardized threshold was used to determine positive staining in each channel. A nuclear neuronal mask was made, consisting of the areas positive for both DAPI and NeuN. Then, an extra-nuclear neuronal mask was constructed, consisting of the areas positive for NeuN but excluding the nuclear neuronal mask, and then contracted by 1 pixel to reduce noise and expanded by 7 pixels to encompass the immediate extracellular space. Both masks were applied to the HMGB1 stain to quantify the fluorescent intensity of HMGB1 in the extranuclear space.

### Volumetric analysis

Brains were sectioned at 30 µm, with every tenth section collected for volumetric analysis. Sections were mounted on microscope slides and allowed to dry before staining for 10 minutes with 1% Sudan Black (Sigma-Aldrich, Cat. #S-2380) made in 70% ethanol (KOPTEC, Cat. #V1016). Next, slides were washed three times with 70% ethanol and three times with PBS and then coverslipped using ProLong Gold Mounting Medium. The stained slices were imaged on a Keyence BZ-X Microscope at x10 magnification. In ImageJ, hippocampal area was measured for each cross-section by outlining the hippocampus. Overall volume was calculated using the following formula: volume = distance between sections (30µm per section x 10 sections in between stained sample sections = 0.3 mm) x sum of areas of all stained sections. Volume was measured across 7 sections, approximately between coordinates AP = -1.2 mm and AP = -3.4 mm.

### Biochemical extraction of brain tissue

Hippocampi previously dissected and stored at -80°C were weighed and combined with ice-cold lysis buffer (RAB buffer; G biosciences, Cat. #786-91) solution at a ratio of 10 µl per mg tissue. Lysis buffer was supplemented with phosphatase inhibitor (Roche, Cat. #PHOSS-RO), cOmplete protease inhibitor (Roche,Cat. #05892791001), nicotinamide (Sigma-Aldrich, Cat. #N0636), and trichostatin A (Abcam, Cat. #ab120850), to prevent various enzymatic breakdowns. Samples were homogenized with handheld homogenizer (Cole-Palmer, Cat. #UX-44468-23) and centrifuged at max speed for 15 minutes at 4°C. Supernatant was saved as brain homogenate. After homogenization, protein levels were determined via Bicinchoninic Acid (BCA) assay. In brief, samples were diluted 10x and incubated with BCA working reagent (50 parts A to 1 part B; Pierce BCA Protein Assay Kit, Cat. #23225) for 30 minutes. Subsequent absorbance was measured with a SpectraMaX M5 spectrophotometer (Molecular Devices) and interpolated against a standard curve to determine protein concentration of each sample.

### Western blot analysis

Hippocampal Lysates were diluted to equal protein concentrations and loaded onto a 12% NuPAGE Bis-Tris polyacrimide gel (Invitrogen, Cat. #NP0343). Proteins were separated via gel electrophoresis at 160V in MOPS running buffer (Invitrogen, Cat. #NP0001) and then transferred onto a nitrocellulose membrane (Bio-Rad, Cat #1620215) at 18V for 60 minutes with a Trans-Blot Turbo Transfer System. Membranes were washed with PBS-T and blocked with intercept blocking buffer for 1 hour (LI-COR, Cat. #927-70001), and then incubated overnight at 4°C with the following primary antibodies: IBA1 (1:1000; Synaptic Systems, Cat. #234308); GFAP (1:1000; Sigma-Aldrich, Cat. #MAB3402); GAPDH (1:2500; Abcam, Cat. #ab83956). Membranes were washed with PBS-T the next day and incubated with the following fluorescently labeled secondary antibodies: Donkey anti-guinea pig 488 (1:2000; Jackson Immuno, Cat. # 706-545-148); Donkey anti-mouse 800RD (1:20000; LI-COR, Cat. #926-32212), and Donkey anti-chicken 680RD (1:20000; LI-COR, Cat. #926-6807) for 1 hour in the dark. Blotted membranes were then scanned using the ChemiDoc MP system (Bio-Rad), and the fluorescent intensity of each band was measured with Image Lab (Bio-Rad). Samples were quantified as the ratio of target protein to protein control: IBA1:GAPDH, and GFAP:GAPDH, normalized to PS19/E4. In some cases, blots were stripped (928-40028, LI-COR) and re-probed for additional antibodies of interest with GAPDH as control.

### Single-nuclei preparation for 10x loading

Single-nuclei preparation was conducted as previously described^14,15^. In brief, hippocampi previously dissected and stored at -80°C were combined with 1 ml of ice-cold homogenization buffer (HB) (250 mM sucrose, 25 mM KCl, 5 mM MgCl2, 20 mM tricine-KOH pH 7.8, 1 mM dithiothreitol, 0.5 mM spermidine, 0.15 mM spermine, 0.3% NP40) supplemented with 0.2 U/µl RNasin Plus RNAse inhibitor (Promega, Cat. #N2615) and protease inhibitor (Roche, Cat. 11836170001). Samples were dounced with ‘A’ loose pestle and ‘B’ tight pestle for 10 strokes each. Homogenate was filtered with a 70 µm Flowmi strainer (Bel-Art, Cat. #H13680-0070) and spun at 350 g for 5 min at 4°C. Supernatant was discarded and the pelleted nuclei were resuspended in 400 μL 1X HB. Next, 400 μl of 50% Iodixanol solution was added to the nuclei to create a 25% mixture, and then slowly layered with 600 μl of 30% Iodixanol solution underneath. Mixture was then layered again with 600 μl of 40% Iodixanol solution under the 30% mixture. The nuclei were spun at 3000 g for 20 min at 4°C before 200 μl of the nuclei band at the 30%–40%interface was combined with 800 μl of 2.5% BSA (Millipore Sigma; Cat. #A9576) in PBS supplemented with RNase inhibitor. Samples were spun for 10 min at 500 g and 4°C. The nuclei were resuspended with 2.5% BSA in PBS supplemented with RNase inhibitor to reach at least 500 nuclei per μl. Nuclei were then filtered with a 40 μm Flowmi stainer (Millipore Sigma; Cat. #BAH136800040) and counted. Roughly 14,000 nuclei per sample were loaded onto a 10x Genomics Next GEM chip G. The snRNAseq libraries were prepared using the Chromium Next GEM Single Cell 3ʹ Library and Gel Bead kit v.3.1 (10x Genomics, Cat. #1000121) according to the manufacturer’s instructions. Libraries were sequenced on an Illumina NovaSeq X Plus sequencer at the UCSF CAT Core.

### Pre-processing and clustering of mouse snRNA-seq samples

The snRNA-seq dataset included 12 total samples, comprising four mice per genotype group (PS19/E3, PS19/E4, PS19/NSE-E4). Each group consisted of 2 male and 2 female mice. The demultiplexed FASTQ files for these samples were aligned to a custom mouse reference genome using the 10x Genomics Cell Ranger Count pipeline^104^ (version 9.0.1), following the Cell Ranger documentation. To accommodate the Homo sapiens MAPT (NCBI reference sequence NM_001123066.4) and the H. sapiens APOE (NCBI reference sequence NM_001123066.4) genes, the custom mouse reference genome was made using the reference mouse genome sequence (GRCm38) from Ensembl (release 98) and the mouse gene annotation file from GENCODE (release M23). The H. sapiens MAPT sequence and H. sapiens APOE sequence were appended as separate chromosomes to the end of the mouse reference genome sequence and the corresponding gene annotations were appended to the filtered mouse reference gene annotation GTF file. The include-introns flag was set to TRUE to capture reads mapping to intronic regions.

The filtered count matrices generated by the Cell Ranger count pipeline for the 12 samples were processed using the R package Seurat^105^ (version 4.4.0) for single-nucleus analysis. Each sample was pre-processed as a Seurat object, then all 12 samples were merged into a single Seurat object. Low-quality nuclei were removed using the following criteria: fewer than 500 total UMI counts, fewer than 200 features, or a mitochondrial gene ratio higher than 0.25%. Sparse gene features expressed in fewer than 10 cells were also excluded. After quality control, normalization and variance stabilization were performed using SCTransform^106^ (version 0.4.1) method for initial parameter estimation. Graph-based clustering was conducted using the Seurat functions, FindNeighbors and FindClusters. Cells were embedded in a k-nearest neighbor (KNN) graph on the first 50 principal components (PCs) using Euclidean distance in the PCA space, and edge weights between two cells were refined using Jaccard similarity. Clustering via the Louvain algorithm (FindClusters) with 15 PCs and a resolution of 0.7 produced 33 distinct biologically relevant clusters, which were subsequently used for further analyses.

### Cell type assignment

Cell type identities were annotated following a method we previously described^107,108^. Briefly, Seurat’s AddModuleScore() function was applied to the object using features from lists of known brain marker genes (up to 8 per cell type)^109,110^. Further subdivisions of hippocampus cell types, such as CA1 and CA3 pyramidal cells, were identified using hippocampus-specific marker genes curated from Hipposeq (https://hipposeq.janelia.org). A module score of each cell type was computed for each individual cell, and each cell was assigned the cell type that produced the highest module score among all tested cell types. Cells with their top two scores within 10% of each other were classified as potential hybrids and excluded from downstream analyses. Cells with negative scores for all cell types computed were classified as negative and excluded from downstream analyses. The cell type assignments per cluster was evaluated by inspecting homogeneity, distribution, and spatial separation of clusters in UMAP space and the expression of major cell type marker genes across clusters. Clusters containing comparable proportions of two or more cell types were deemed unknown or mixed.

### Gene set enrichment analysis

DE genes between clusters of interest or between two genotype groups were identified using the FindMarkers Seurat function on the SCT assay data. This algorithm uses the Wilcoxon rank-sum test to identify DE genes between two populations. DE genes were limited to genes detected in at least 10% of the cells in either population and with at least 0.1 log_2_ fold change. Volcano plots with log_2_ fold change and p value from the DE gene lists were generated using the ggplot2 R package^111^ (version 3.5.2). Overrepresentation (or enrichment) analysis was performed using clusterProfiler^112^ (version 4.14.6) to find gene sets associated with the DE genes with at least 10 genes in the mouse-specific KEGG database^113^. The p values are based on a hypergeometric test and are adjusted for multiple testing using the Benjamini–Hochberg method^114^.

### Association between clusters and genotype

A GLMM_AM was implemented in the lme4 (version 1.1-37) R package^115^ and used to estimate the associations between cluster membership and the mouse model. These models were run separately for each cluster of cells. The GLM model was performed with the family argument set to the binomial probability distribution and with the ‘nAGQ’ parameter set to 10, corresponding to the number of points per axis for evaluating the adaptive Gauss–Hermite approximation for the log-likelihood estimation. Cluster membership of cells by sample was modeled as a response variable by a two-dimensional vector representing the number of cells from the given sample belonging to and not belonging to the cluster under consideration. The corresponding mouse ID from which the cell was derived was the random effect variable, and the animal model for this mouse ID was included as the fixed variable. The reference animal model was set to PS19/E3. The resulting p values for the estimated LOR across the three animal models (with respect to the PS19/E3) and clusters were adjusted for multiple testing using the Benjamini–Hochberg method^114^.

### Association between proportion of cell types and histopathological parameters

A GLMM_histopathology was implemented in the lme4 (version 1.1-37) R package^115^ and used to identify cell types whose proportions are significantly associated with changes in histopathology across the samples. These models were performed separately for each combination of all clusters and each histological parameter. The GLM model was performed with the family argument set to the binomial probability distribution family and with the ‘nAGQ’ parameter set to 1 corresponding to a Laplace approximation for the log-likelihood estimation. Cluster membership of cells by sample was modeled as a response variable by a two-dimensional vector representing the number of cells from the given sample belonging to and not belonging to the cluster under consideration. The corresponding mouse model from which the cell was derived was included as a random effect, and, further, the mouse ID within the given mouse model was modeled as a random effect as well. Note, this represents the hierarchical nature of these data for the GLMM, and the mouse models are first assumed to be sampled from a ‘universe’ of mouse models; this is then followed by sampling mice within each mouse model. The modeling choice of including the mouse model as a random effect as opposed to a fixed effect is meant to increase the degrees of freedom (or maximize the statistical power) to detect the association of interest, particularly in light of the relatively small number of replicates (3–4) per animal model. The histological parameter under consideration was modeled as a fixed effect in this model.

We visualized the LOR estimates (derived from the GLMM fits) in a heatmap using pheatmap package 1.0.12 after adjusting the p values distribution across histopathological parameters across cell types with Benjamini–Hochberg multiple testing correction^114^.

### General Statistics and Reproducibility

Sample sizes for mouse studies were chosen on the basis of estimates to provide statistical power of ≥80% and alpha of 0.05 based on preliminary data. The differences between genotype groups were evaluated by ordinary one-way ANOVA with Tukey’s multiple comparisons test, where the mean of each column was compared to the mean of every other column. All data are shown as mean ± s.e.m. Data distribution was assumed to be normal, though this was not formally tested. For correlations between two data sets, simple linear regression was used. p < 0.05 was considered to be significant, and all significant p values were included in figures or noted in figure legends. Statistical significance analysis and plots were completed with GraphPad Prism 10 for Mac (GraphPad Software). No randomization method was used for the assignment of mice to study groups, and no animals or data points were excluded from these studies. For mouse snRNA-seq studies, sample sizes were determined by a power analysis based on effect sizes from our previous studies and literature^14,15^. Mice selected for snRNA-seq consisted of 2 males and 2 females per group that together represented average pathologies for all parameters quantified. Nuclei were isolated from four mice per genotype to ensure n ≥ 3 mice per group. Pathology data were correlated with snRNA-seq data. Investigators were not blinded during analysis of the snRNA-seq datasets, as sample metadata were needed to conduct any comparisons. Studies were all performed using one cohort of mice.

## Data availability

The H. sapiens MAPT sequence is available at https://www.ncbi.nlm.nih.gov/nuccore/NM_001123066. The reference mouse genome sequence (GRCm38) from Ensembl (release 98) is available at http://ftp.ensembl.org/pub/release-98/fasta/mus_musculus/dna/Musmusculus.GRCm38.dna.primary_assembly.fa.gz. The reference mouse gene annotation file from GENCODE (release M23) is available at http://ftp.ebi.ac.uk/pub/databases/gencode/Gencode_mouse/release_M23/gencode.vM23.primary_asse mbly.annotation.gtf.gz. The snRNA-seq datasets generated during the study are available at the Gene Expression Omnibus under accession number (link).

Data associated with Figs. 6 and 7 and Extended Data Figs. 5–7 are also available in the Supplementary Information. Source data are provided with this paper.

## Code availability

The following packages/software were used either as dependencies to downloading or using packages mentioned in the Methods section or in creating the figures in this manuscript: Seurat v.4.4.0, CellRanger v.9.0.1, glm lme4 v.1.1-37, clusterProfiler v.4.14.6, ggplot2 v.3.5.2, pheatmap v.1.0.12, and Fiji v.2.1.0 (ImageJ). Western blot images were analyzed using Image Lab v.6.1.0. GraphPad Prism v.10.6.1 was used for some statistical analyses and graphing. All codes generated with custom R and shell scripting during this study are accessible via GitHub (link).

## Acknowledgements

This work was partially supported by National Institutes of Health (NIH) grants R10AG071697, RF1AG076647, and P01AG073082 to Y. Huang and the UCSF ARCS 2022-2023 and 2023-2024 Fellowships awarded to J. B. The funders had no role in study design, data collection and analysis, decision to publish or preparation of the manuscript. We thank the entire Huang laboratory and Gladstone Institute of Neurological Disease colleagues for invaluable discussions and input; R. Blandino of the Gladstone Histology and Light Microscopy Core for microscopy training; the UCSF Cat core, for sequencing, supported by UCSF PBBR, RRP IMIA, and NIH 1S10OD028511-01 grants; and T. Pak for editorial assistance.

## Author Contributions

J.B. and Y. Huang collaboratively designed the experiments and wrote the manuscript. J.B. performed the majority of studies, data collection, and analyses. Y.L. and L.Y. performed the snRNA-seq analyses and generated sequencing data plots. M.J.K. aided in tissue collection and processing. M.J.K., O.Y., R.F., D.S., and Z.P. assisted in some immunohistochemical staining and microscopy. R.F. conducted some western blot experiments. N.K. shared some immunohistochemical protocols and tissue. K.S. and Y.Hao isolated cell nuclei and prepared samples for snRNA-seq. C.E., S.D.L., and J.N. managed all mouse lines. S.D.L coordinated all genotyping. Y.Huang supervised the project.

---

**Extended Data Fig. 1.**
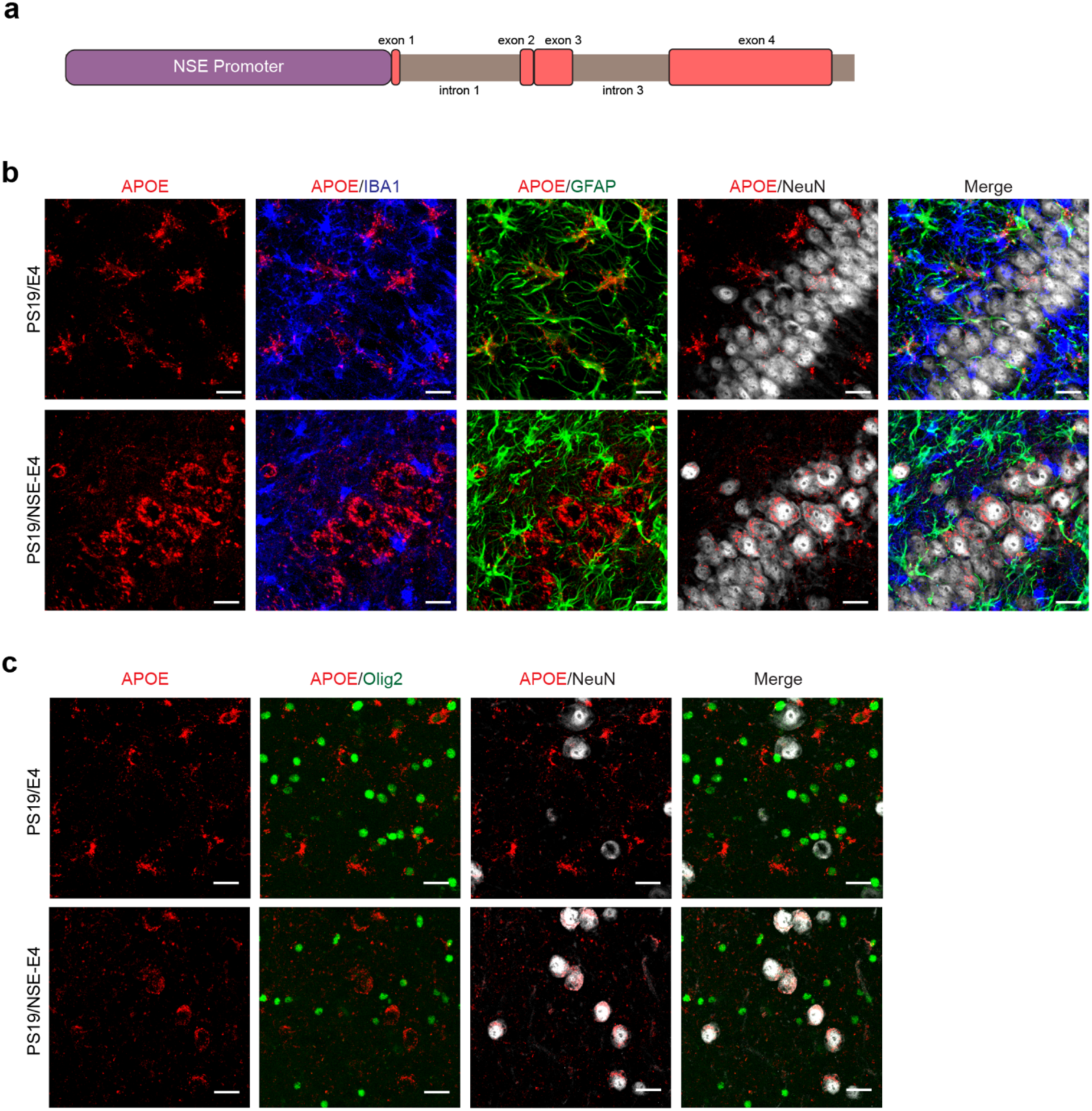
Schematic of APOE4 minigene construct and validation of the PS19/NSE-E4 mouse model with APOE selectively expressed by neurons. **a**, Schematic of APOE4 minigene inserted under the NSE promoter. **b,c**, Representative images of the cell-type specificity of APOE expression in the hippocampus of 10-month-old PS19/E4 and PS19/NSE-E4 mice. In **b,** APOE is depicted in red, co-labeled with microglia (IBA1) in blue, astrocytes (GFAP) in green, and neurons (NeuN) in white (scale bar, 20 µm). In **c,** APOE is depicted in red, co-labeled with oligodendocytes (Olig2) in green and neurons (NeuN) in white (scale bar, 20 µm).

**Extended Data Fig. 2.**
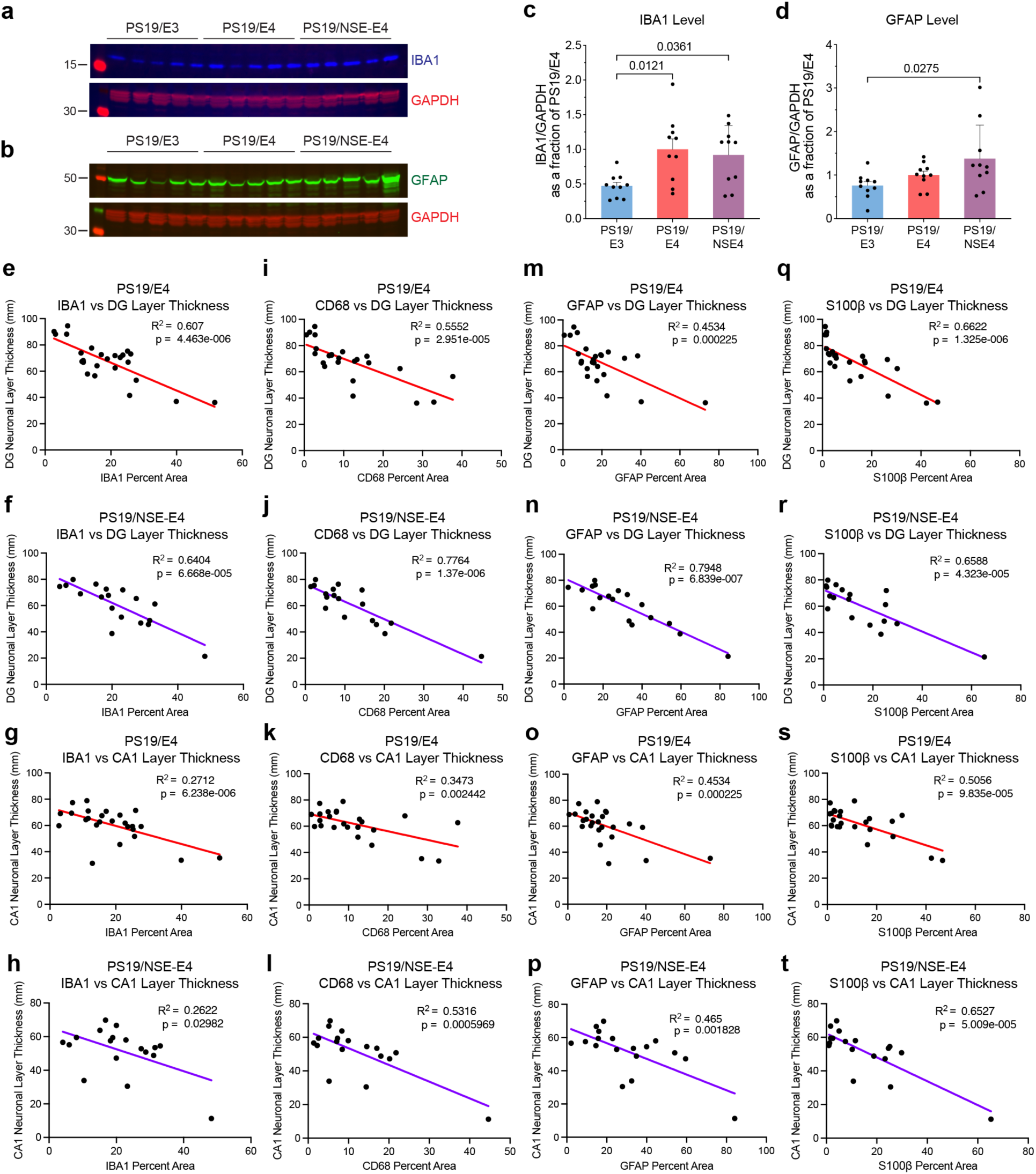
Neuronal APOE4-promoted gliosis is correlated with neurodegeneration. **a**, Representative western blot images with microglia IBA1 antibody. GAPDH served as a loading control. **b**, Representative western blot images with astrocyte GFAP antibody. Membrane in **a** was stripped and stained for GFAP and GAPDH. GAPDH served as a loading control. **c,d**, Quantifications of IBA1 (**c**) and GFAP (**d**) levels in hippocampal lysates of PS19/E3 (n = 10), PS19/E4 (n = 10), and PS19/NSE-E4 (n = 10) mice. Immunoblot levels were normalized to GAPDH first and then to those of PS19/E4 mice. **e,f**, Correlation of IBA1 microglia levels and DG layer thickness in PS19/E4 mice (**e**) and PS19/NSE-E4 mice (**f**). **g,h**, Correlation of IBA1 microglia levels and CA1 layer thickness in PS19/E4 mice (**g**) and PS19/NSE-E4 mice (**h**). **i,j**, Correlation of CD68 microglia levels and DG layer thickness in PS19/E4 mice (**i**) and PS19/NSE-E4 mice (**j**). **k,l**, Correlation of CD68 microglia levels and CA1 layer thickness in PS19/E4 mice (**k**) and PS19/NSE-E4 mice (**l**). **m,n**, Correlation of GFAP astrocyte levels and DG layer thickness in PS19/E4 mice (**m**) and PS19/NSE-E4 mice (**n**). **o,p**, Correlation of GFAP astrocyte levels and CA1 layer thickness in PS19/E4 mice (**o**) and PS19/NSE-E4 mice (**p**). **q,r**, Correlation of S100β astrocyte levels and DG layer thickness in PS19/E4 mice (**q**) and PS19/NSE-E4 mice (**r**). **s,t**, Correlation of S100β astrocyte levels and CA1 layer thickness in PS19/E4 mice (**s**) and PS19/NSE-E4 mice (**t**). In **f**-**t**, PS19/E3, *n* = 24; PS19/E4, *n* = 25; and PS19/NSE-E4, *n* = 18. Correlations utilized Pearson correlation analysis to determine significance.

**Extended Data Fig. 3.**
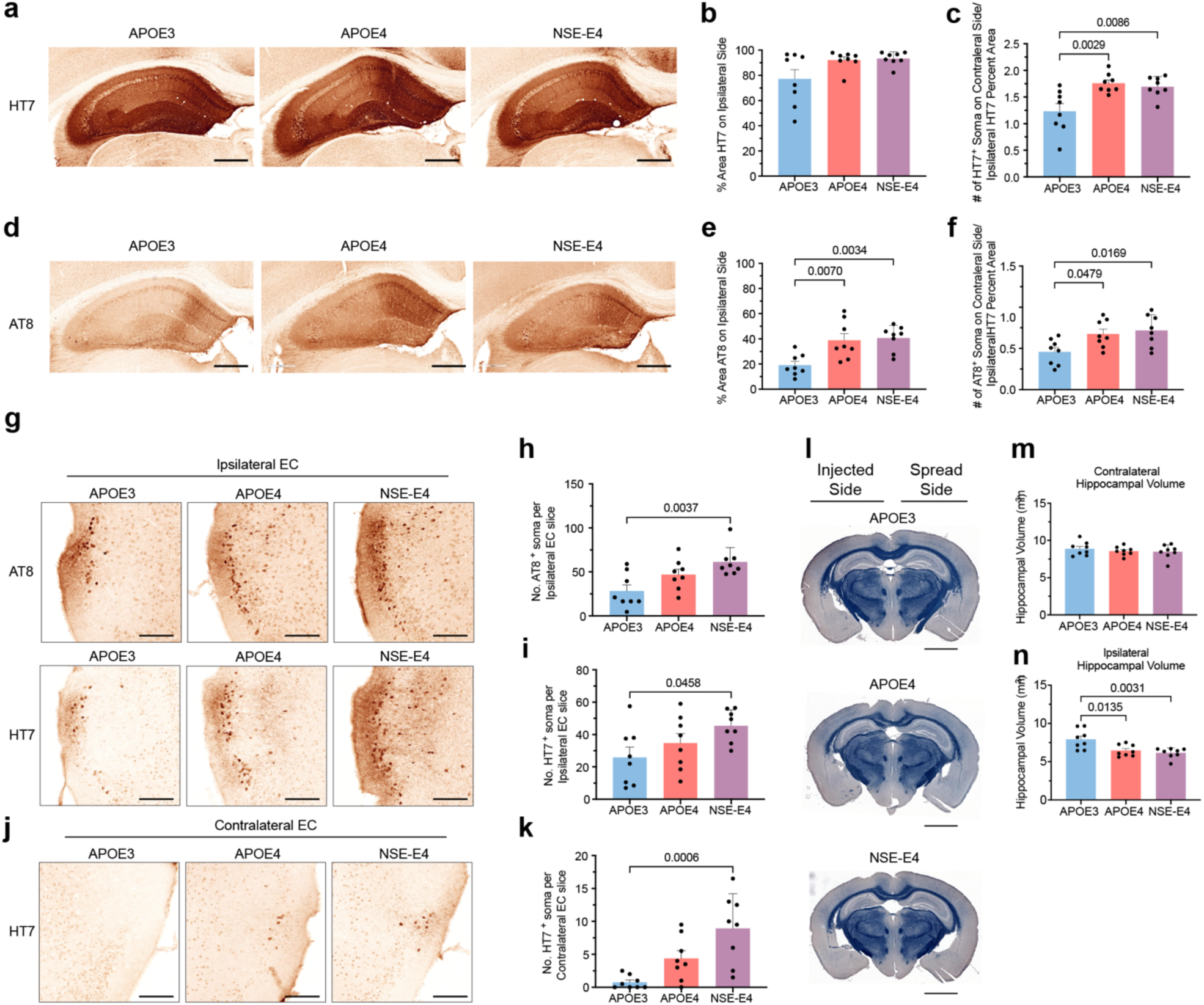
Neuronal APOE4 alone drives additional pathological endpoints in the *in vivo* Tau propagation model. **a**, Representative images of human Tau (HT7) immunolabeling of the ipsilateral hippocampus of APOE3, APOE4, and NSE-E4 mice, 10 weeks post unilateral injection of mutant human Tau virus (scale bar, 500 µm). **b**, Quantifications of the percent HT7 coverage area in the ipsilateral hippocampus of these mice. **c**, Quantifications of the number of HT7-positive soma on the contralateral side normalized to the percent HT7 coverage area of the ipsilateral side of these mice. **d**, Representative images of p-Tau (AT8) immunolabeling of the ipsilateral hippocampus of APOE3, APOE4, and NSE-E4 mice, 10 weeks post unilateral injection of mutant human Tau virus (scale bar, 500 µm). **e**, Quantifications of the percent AT8 coverage area in the ipsilateral hippocampus of these mice. **f**, Quantifications of the number of AT8-positive soma on the contralateral side normalized to the HT7 percent coverage area of the ipsilateral side of these mice. **g**, Representative images of human Tau (HT7) and p-Tau (AT8) immunolabeling of the ipsilateral entorhinal cortex (EC) of injected APOE3, APOE4, and NSE-E4 mice (scale bar, 200 µm). **h,i**, Quantifications of the number of AT8-positive cells (**h**) and HT7-positive cells (**i**) in the ipsilateral EC of these mice. **j**, Representative images of human Tau (HT7) immunolabeling of the contralateral EC of injected APOE3, APOE4, and NSE-E4 mice (scale bar, 200 µm). **k**, Quantifications of the number of HT7-positive cells in the contralateral EC of these mice. **l**, Representative hippocampal cross-sections of injected APOE3, APOE4, and NSE-E4 mice, stained with Sudan Black to enhance hippocampal visualization (scale bar, 2000 µm). **m,n**, Quantifications of the contralateral (**m**) and ipsilateral (**n**) hippocampal volumes of these mice. For quantifications in **b**,**c**,**e**,**f**,**h**,**i**,**k**,**m**,**n**, APOE3 *n* = 8; APOE4 *n* = 8; and NSE-E4 *n* = 8; data is expressed as mean ± s.e.m and was assessed via one-way analysis of variance (ANOVA) with Tukey’s post hoc multiple comparisons test.

**Extended Data Fig. 4.**
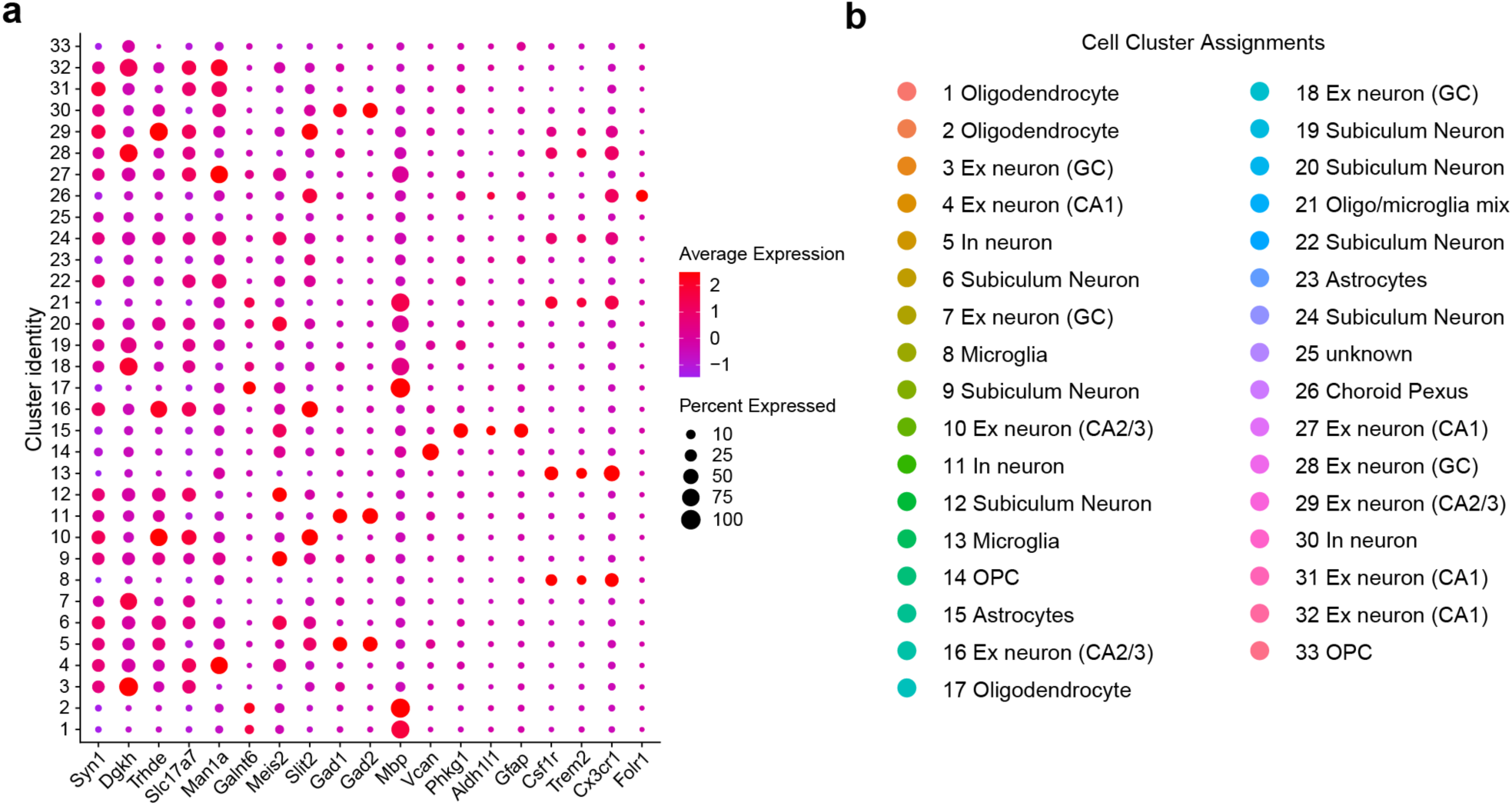
snRNA-seq identification of cell clusters. **a**, Dot-plot depicting normalized average expression of selected cell identity marker genes for all 33 unique hippocampal cell clusters from mice at 10 months of age. The size of the dots is proportional to the percentage of cells expressing a given gene. Average expression is on a colored scale (lower expression, blue; higher expression, red). **b**, Assigned identity of 33 distinct cell types. In neuron, inhibitory neuron; Ex neuron, excitatory neuron; oligo, oligodendrocyte.

**Extended Data Fig. 5.**
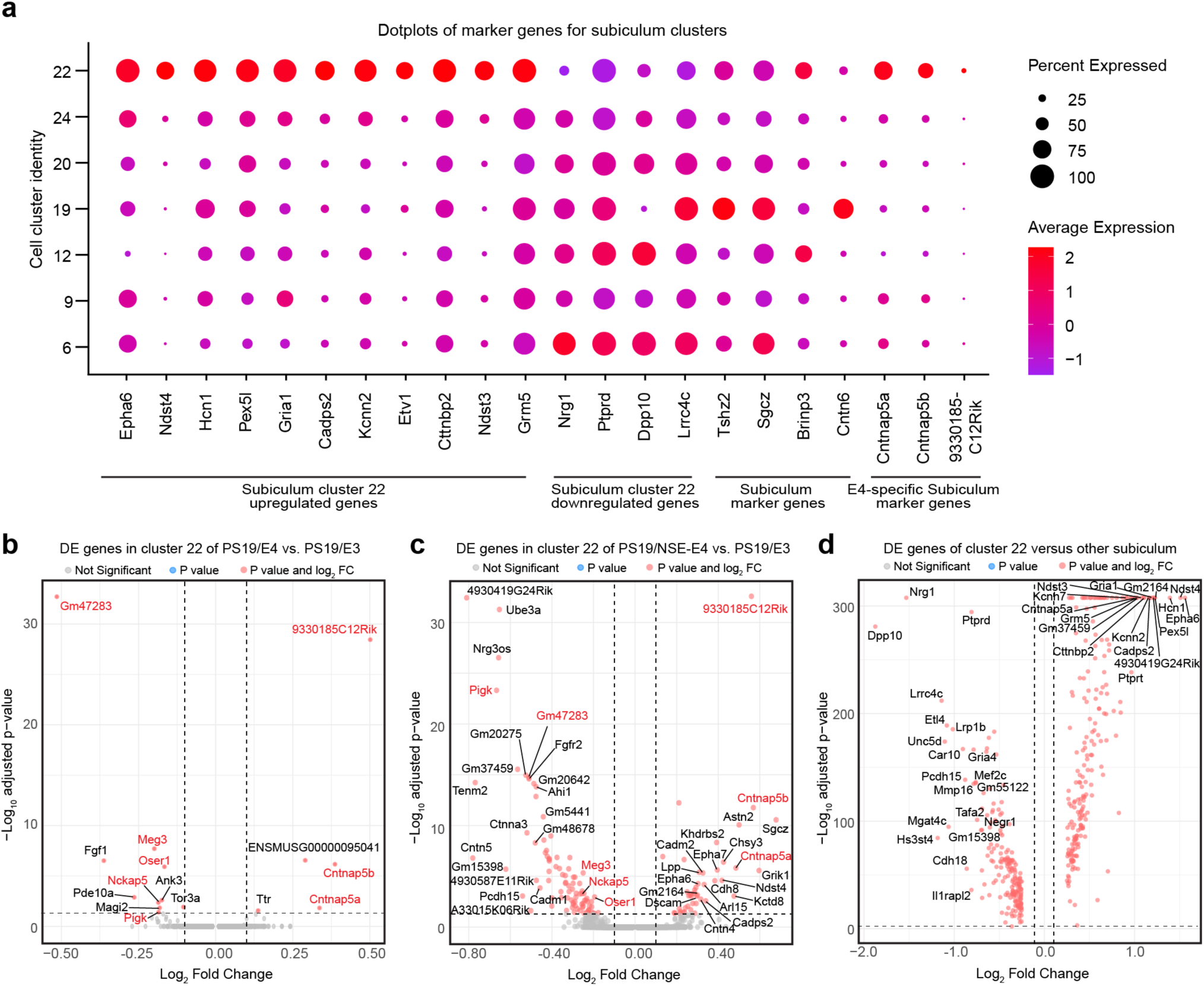
Characteristics of neuronal APOE4-vulnerable subiculum neuron cluster 22. **a**, Dot-plot of normalized average expression of marker genes and genes of interest for selected subiculum neuron clusters. The size of the dots is proportional to the percentage of cells expressing a given gene. Average expression is on a colored scale (lower expression, blue; higher expression, red). **b**, Volcano plot of the differentially-expressed (DE) genes between PS19/E4 and PS19/E3 in cluster 22 subiculum neurons. **c**, Volcano plot of the DE genes between PS19/NSE-E4 and PS19/E3 in cluster 22 subiculum neurons. Up- or down-regulated genes shared between **b** and **c** are indicated in red font. **d,** Volcano plot of the DE genes between subiculum neuron cluster 22 and all other subiculum clusters. For all volcano plots in **b**-**d**, Dashed lines represent log_2_ fold change threshold of 0.1 and p value threshold of 0.05. The unadjusted p values and log_2_ fold change values used were generated from the gene-set enrichment analysis using the two-sided Wilcoxon rank-sum test as implemented in the FindMarkers function of the Seurat package.

**Extended Data Fig. 6.**
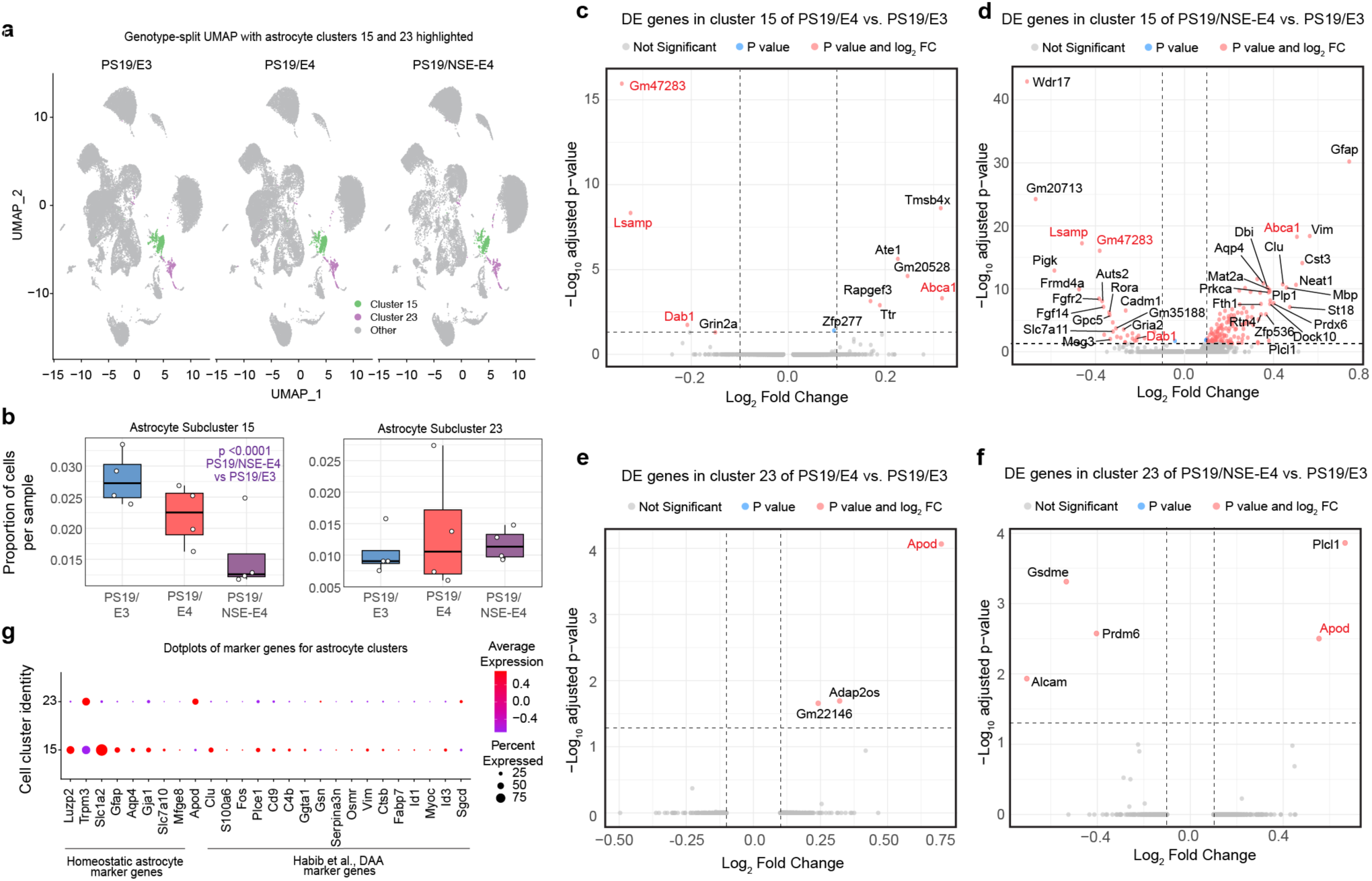
Neuronal APOE4 induces mild astrocyte changes. **a**, Genotype-split UMAP highlighting astrocyte clusters 15 and 23 across each genotype group. **b**, Box plot of the proportion of cells from each sample for astrocyte clusters 15 and 23 (PS19/E3, *n* = 4; PS19/E4, *n* = 4; and PS19/NSE-E4, *n* = 4). From bottom to top, the hinges of the box plots correspond to the 25^th^, 50^th^, and 75^th^ percentiles. The upper and lower whiskers of the box plot extend to the largest and smallest values, respectively, though no further than 1.5 x IQR from the nearest hinge. IQR, interquartile range, or distance between the 25^th^ and 75^th^ percentiles. The log odds ratios (LOR) are the mean ± s.e.m estimates of LOR for these clusters, which represents the change in the log odds of cells per sample from each mouse belonging to the respective clusters relative to log odds of cells per sample from PS19/E3 mice. **c**, Volcano plot of the differentially-expressed (DE) genes between PS19/E4 and PS19/E3 in cluster 15 astrocytes. **d**, Volcano plot of the DE genes between PS19/NSE-E4 and PS19/E3 in cluster 15 astrocytes. Up- or down-regulated genes shared between **c** and **d** are indicated in red font. **e**, Volcano plot of the DE genes between PS19/E4 and PS19/E3 in cluster 23 astrocytes. **f**, Volcano plot of the DE genes between PS19/NSE-E4 and PS19/E3 in cluster 23 astrocytes. Up- or down-regulated genes shared between **e** and **f** are indicated in red font. **g**, Dot-plot of normalized average expression of marker genes and genes of interest for astrocyte clusters. The size of the dots is proportional to the percentage of cells expressing a given gene. Average expression is on a colored scale (lower expression, blue; higher expression, red). Unadjusted p values in **b** are from fits to a GLMM_AM; association tests were two-sided. In **c**-**f**, dashed lines represent log_2_ fold change threshold of 0.1 and p value threshold of 0.05; the unadjusted p values and log_2_ fold change values used were generated from the differential expression analysis using the two-sided Wilcoxon rank-sum test as implemented in the FindMarkers function of the Seurat package.

